# Simultaneous profiling of spatial gene expression and chromatin accessibility for mouse brain development

**DOI:** 10.1101/2022.03.22.485333

**Authors:** Fuqing Jiang, Xin Zhou, Yingying Qian, Miao Zhu, Li Wang, Zhuxia Li, Qingmei Shen, Fangfang Qu, Guizhong Cui, Kai Chen, Guangdun Peng

## Abstract

Brain are complex biological tissues which function relies on coordinated anatomical and molecular structure comprised by a large number of specialized cells. The spatial architecture of brain which is key to the understanding of its physiological and pathological significance is formed during embryo development. However, the molecular basis for discrete neuroanatomical domains particularly in the context of spatial organization of the brain is inadequate. Here, we introduced microfluidic indexing based spatial ATAC and RNA sequencing (MISAR-seq), a method for joint profiling of chromatin accessibility and gene expression with spatial information retained in the developing mouse brain. Our study has established a direct means to spatially determine the coordination between chromatin accessibility and transcriptome, identified the chromatin potential to define cell fate determination of brain organization, and uncovered spatiotemporal regulatory principles during mammalian brain development.

---

The brain is an enormously complex organ that composed of spatially organized and functionally interconnected neurons, glia and non-neural cells. An entry point in understanding brain development and function is to annotate the atlas of the brain by highly scalable and unbiased single-cell multi-omics approaches^1–4^. In recent years, brain region formation and cell maturation process have been profiled using a variety of single cell technologies, including single cell RNA sequencing (scRNA-seq)^2^, single nuclear ATAC sequencing (snATAC-seq)^5, 6^, single cell multimodal technologies^7^ and single cell joint multi-omics sequencing^3^, which uncovered the complex interplay of brain neurons cell-cell interaction and led to the deep understanding of intrinsic genetic programs of multiple cell types. With the advent of spatial transcriptome technologies^8, 9^, the molecular cytoarchitecture of the brain has been unveiled in unprecedented detail. A complete description on spatially resolved transcriptome atlas for the developing and adult brain is on the horizon. However, one big challenge to define the building blocks of brain function lies in deciphering the complex regulatory network, e.g. the connection between chromatin modification and its transcription output, in forming the sophisticated tissue organization.

Similar to the advances in single-cell sequencing, spatial multiomics is the new frontier for linking measurements from layers of genome, epigenome, transcriptome and proteomes at the spatial level to reveal regulatory and functional mechanisms. In the past few years, there has been a rapid emergence of spatial transcriptome technologies on different platforms, together with the infant spatial epigenome, greatly expanding the toolkits for our understanding the interplay of biological networks. Importantly, different modalities of spatial dimension (e.g., spatial-ATAC-seq^10^, spatial-cut&tag^11^) can be achieved individually, there is a requisite to simultaneously profile the gene expression and regulatory genomic information in the spatial context on a large scale, by which insightful gene regulatory interactions can be gained and is essential for dissecting the basis of anatomical tissue architecture. Here, we reported a spatially resolved joint profiling of chromatin accessibility and gene expressions approach (MISAR-seq) and we applied it to mouse brain of consecutive developmental stages. We showed that high quality transcriptome-open chromatin status in the different anatomical brain regions can be delineated and our data revealed the dynamic spatiotemporal regulatory mechanisms in establishing the complex architecture of mouse brain.

## Results

### Overview of MISAR-seq design and data quality

We devised a microfluidic indexing based spatial ATAC and RNA sequencing (MISAR-seq) method, motivated by DBiT-seq design^12^ and SHARE-seq methodology^13^ (Extended Data Fig. 1a-c). MISAR-seq includes the following steps (Extended Data Fig. 1a). First, we modified the flow of microchannels and sealing slabs to facilitate a leakage-prevent and minimized diffusion effect (Extended Data Fig. 1b,c). After fixed by 1% formaldehyde, Tn5 transposase assembled with read1 and read2 adaptor sequence is administrated to the tissue slide to label the open chromatin regions in situ. The tissue slide is refixed and mRNA is reverse transcribed using a poly (T) primer bearing an 10bp unique molecular identifier (UMI), read2 anchor sequence and a biotin tag. Next, the tissue section is put onto the microfluidic chip and a set of DNA barcode round1_i (i = 1 to 50) are flowed over the tissue surface through microchannel-guided delivery to ligate to the read2 anchors. A second set of DNA barcode round2_i are flowed over the same tissue surface through microchannels perpendicular to the tissue slides and are attached to the ends of the barcode 1. These two rounds of ligation ensure a spatial barcoding to paired ATAC and mRNA molecules of at most 2500 grids (currently 50 μm in width, representing around 15-30 cells). Adjacent sections are stained to obtain a high-sensitivity anatomical image for downstream alignment. As a validation of leakage-free, Cy3 and Fam dyes are added with the two rounds of barcodes respectively. Additionally, these two dyes can form a 2D mosaic pattern on the tissue slides for coordinates calibration aided with hematoxylin and eosin (H&E) images (Extended Data Fig. 1d). Typically, we would get nucleosomal distribution library for ATAC, no less than 800 bp amplicons and 300∼700 bp cDNA fragments for RNA library at the end of the experimental (Extended Data Fig. 1e-g).

**Fig. 1.**
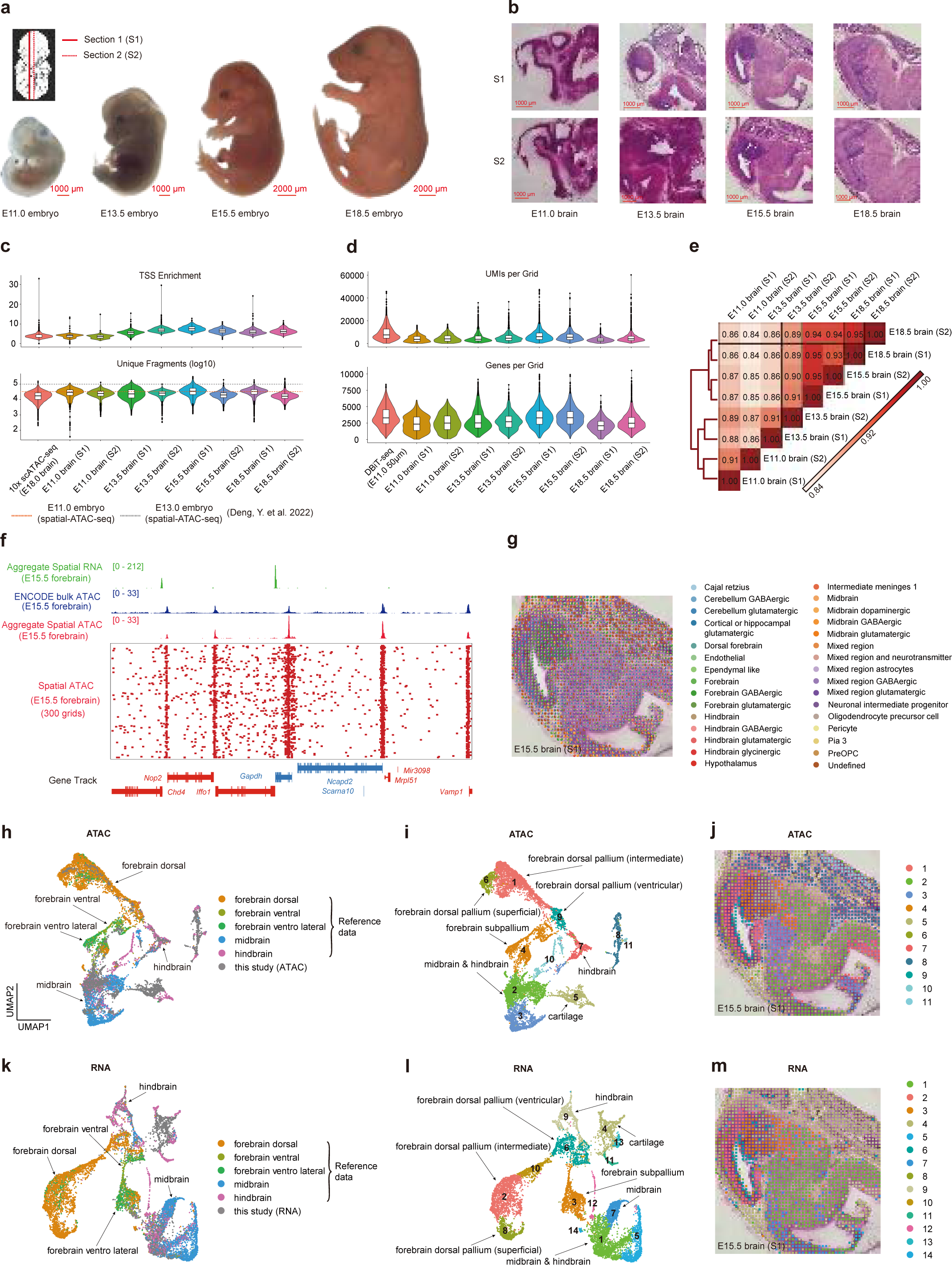
Overview of MISAR-seq: data quality and validation. **a**, Overview of the sampled embryonic time points. **b**, H&E images from an adjacent tissue section. **c,** Comparison of TSS Enrichment value (top) and unique fragments (bottom) per grid between MISAR-seq and 10x scATAC-seq. Dashed lines indicate the results from Spatial-ATAC-seq^10^.. **d**, Comparison of UMIs (top) and genes number (bottom) per grid between MISAR-seq and DBiT-seq. **e**, Spearman correlation coefficient of bulk profiles from eight embryo mouse brain sections in this experiment. **f**, Aggregate selected forebrain spatial RNA and ATAC peak profile compared to published profiles of ATAC-seq in E15.5 mouse forebrain. **g**, Spatial distribution of E15.5 brain spatial RNA single cell deconvolution results with E15.5 brain scRNA-seq reference data. **h**-**j**, The clustering results of E15.5 mouse brain for spatial ATAC after integration with reference E15.5 brain scRNA-seq data. UMAP results annotated with reference (h) and anatomic (i) or spatial mapping (j). **k-m**, The clustering results of E15.5 mouse brain for spatial RNA after integration with reference E15.5 brain scRNA-seq data. UMAP results annotated with reference (k) and anatomic (l) or spatial mapping (m).

We performed MISAR-seq on the fetal brain of E11.0, E13.5, E15.5 and E18.5 mouse embryo (Fig. 1a,b). For spatial ATAC of MISAR-seq, the TSS enrichment score calculated among the unique nuclear fragments showed higher coverage of TSSs, compared to 10x scATAC-seq data (Fig 1c) . A median of 16,472 to 34,730 unique fragments per grid was achieved after filtering low quality reads, of which between 15.0% to 45.6% fragments fell within peak regions (FRiP), indicating a high coverage and low background. In comparison, the unique fragments per cell from 10x scATAC-seq of E18.0 brain was 17,368 and the FRiP was 24.7%. A recently reported spatial-ATAC-seq based on similar platform had, on average, 36,303 unique fragments for E11.0 embryo and 100786 for E13.0 embryo while the FRIP value was 10.0% for E11.0 embryo and 8.0% for E13.0 embryo per gird in the same spatial resolution^11^ (Fig. 1c and Extended Data Fig. 2a). The TSSs enrichment score calculated among the unique fragments showed higher coverage of TSSs, compared to 10x scATAC-seq and recent spatial-ATAC-seq data (Extended Data Fig. 2a). The fractions of read-pairs mapped to mitochondrial of each sample section were between 2.52% to 2.95%, which were slightly higher than 10x scATAC-seq platform but is reasonable due to the openness of the cells on tissue sections (Extended Data Fig. 2a). Meanwhile, our MISAR-seq showed typical nucleosomal and subnucleosomal fragments distribution (Extended Data Fig. 1e, 2b) and captured more transcription start sites (TSSs) compared to 10x scATAC-seq (Fig. 1c and Extended Data Fig. 2c and 3).

**Fig. 2.**
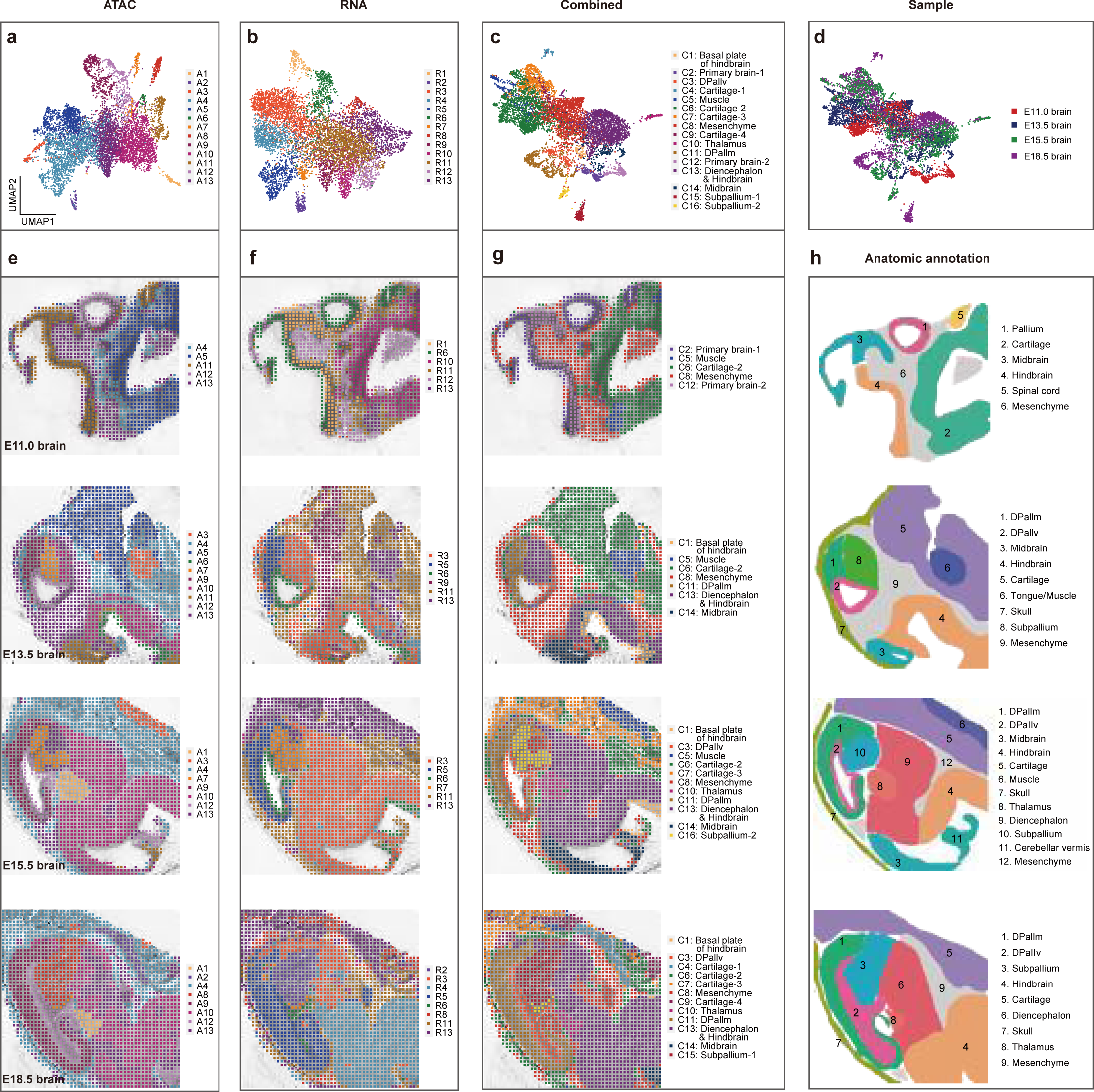
Spatial chromatin accessibility, gene expression and combined mapping for mouse brain development at E11.0, E13.5, E15.5 and E18.5. a-c**, Spatial ATAC (a), RNA** (b) and combined (c) UMAP visualization of different mouse brain development stage, colored by different clusters. **d**, Combined spatial ATAC and RNA UMAP visualization of integrated different mouse brain development stage, colored by different sample sections. **e-g**, Unsupervised clustering of mouse brain sections for spatial ATAC (e), RNA (f) and combined (g). **h**, Anatomic annotation of major tissue regions based on the H&E images for different mouse brain stages. DPallm, mantle zone of dorsal pallium. DPallv, ventricular zone of dorsal pallium.

**Fig. 3.**
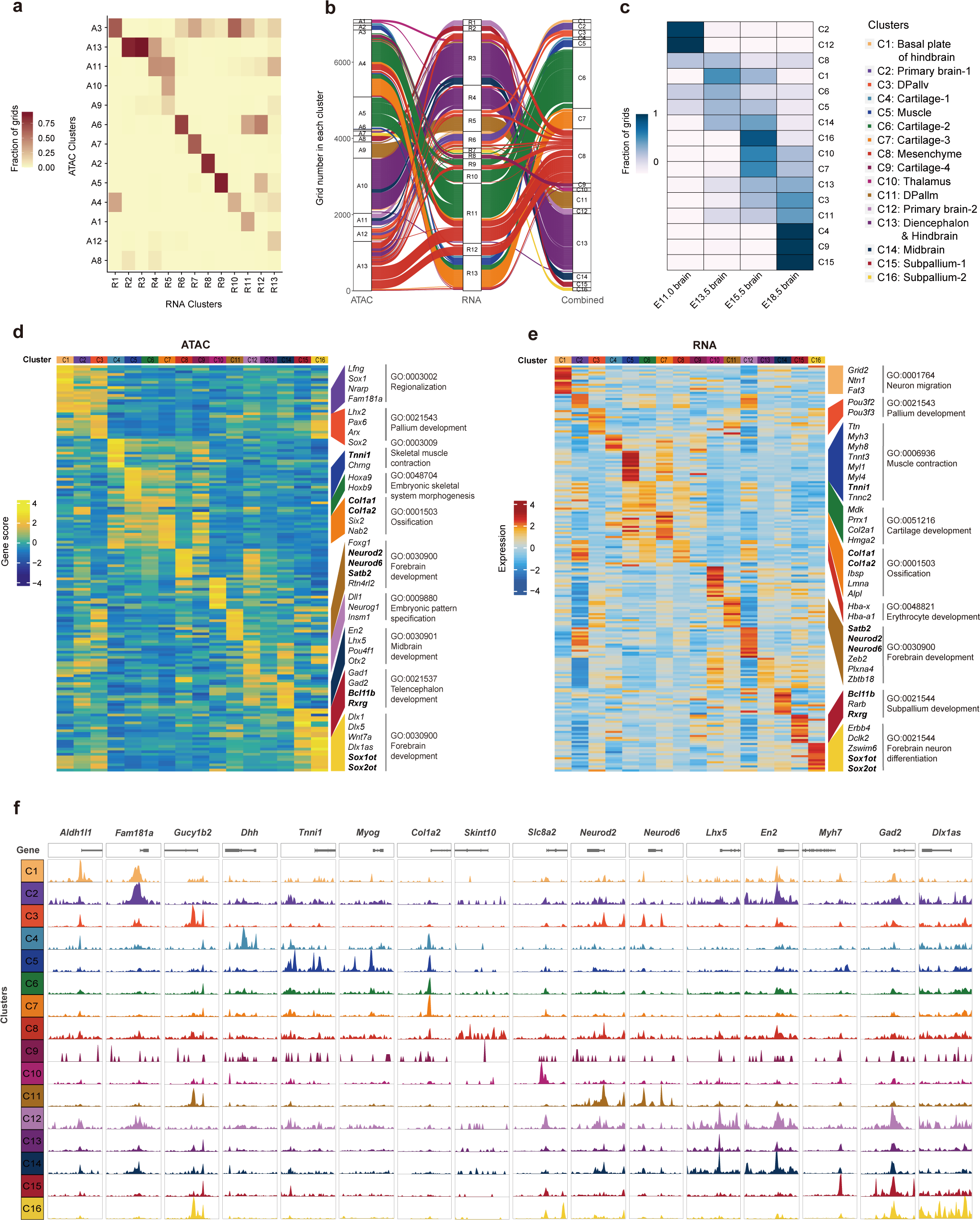
Combined analysis of chromatin accessibility and gene expression across development stages of mouse brain. **a**, Heatmap showing the proportion of cells in the ATAC clusters that overlap with RNA clusters. **b**, Cluster connective changes between ATAC, RNA and combined clusters shown in sankey plots. **c**, The proportion of cells in the combined clusters for development stages of mouse brain. **d-e**, Heatmap of spatial ATAC (d) and RNA (e) marker genes across all clusters. Select GO terms and top ranked genes are shown. **f**, Representative genome browser views of marker genes from aggregated spatial ATAC data from MISAR-seq.

To further validate the ATAC data of MISAR-seq, we compared the MISAR-seq ATAC signals with bulk ATAC data from ENCODE at the same E15.5 brain tissue^14^. The results showed that they shared major ATAC peaks in all forebrain, midbrain and hindbrain regions (Extended Data Fig. 4a-c). We also calculated the overlapped peaks from aggregated brain regions and found most of the peaks in the MISAR-seq ATAC study were also found in the Encode bulk data (Extended Data Fig. 4d-f).

**Fig. 4.**
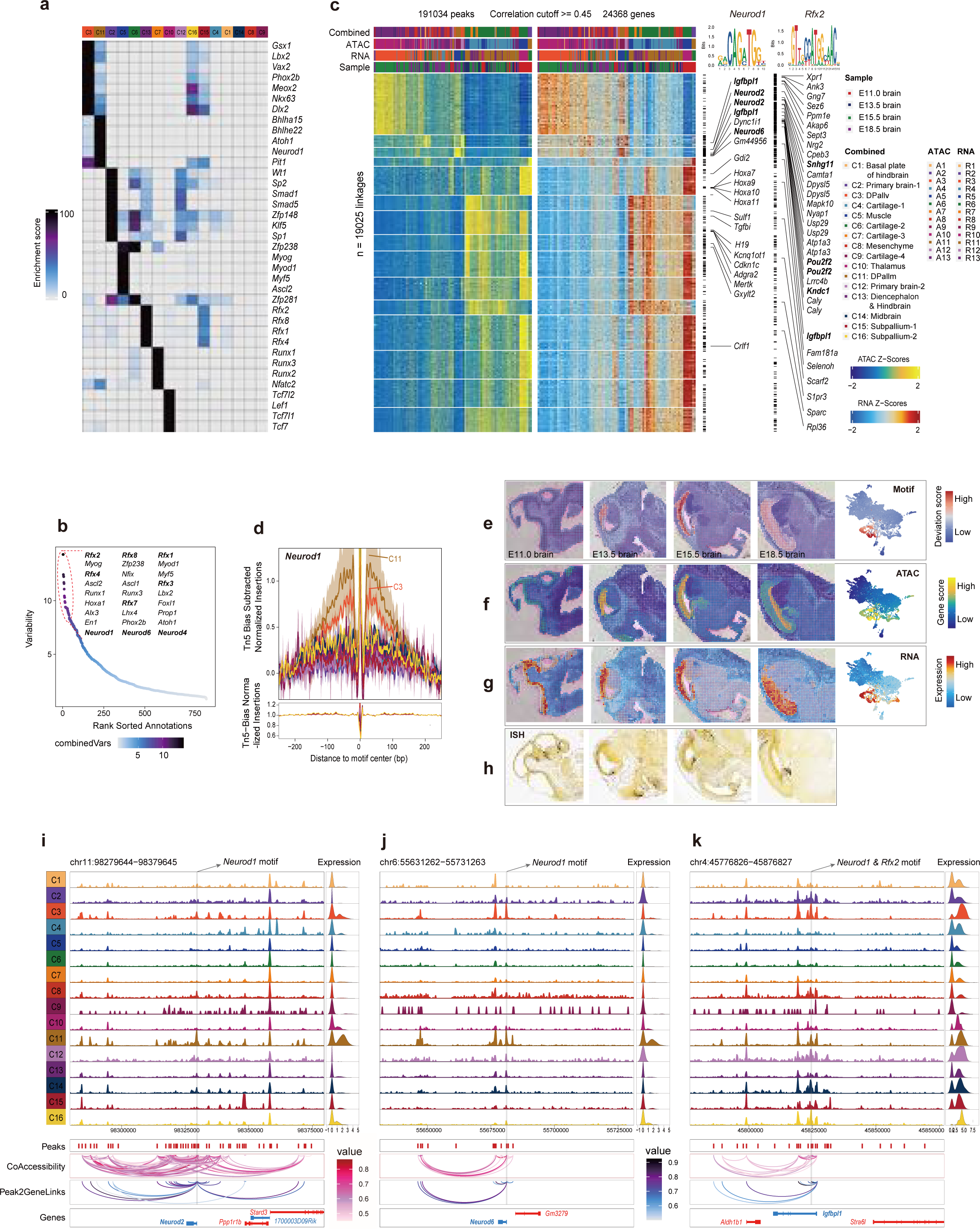
The key role of *Neurod1* and *Rfx2* regulator in mouse brain development. **a**, Enriched marker TFs for each cluster. **b**, Ranked TFs based on the deviation score and top TFs were listed. **c**, Heatmap showing chromatin accessibility and gene expression of 11772 significantly peak-gene linkages. Shown are side-by-side heatmaps in which one row represents a pair of one peak and one linked gene. Each peak can be linked to multiple genes, and each gene can be linked to multiple peaks. the peaks that correlation >=0.7 in peak-gene linkages contained motif of *Neurod1* and *Rfx2* were labeled at right. **d**, Tn5 bias-adjusted transcription factor footprints for *Neurod1* motifs. **e-g**, UMAP embedding and spatial mapping of gene scores (e), gene expression (f), TF deviation score and in situ hybridization results (g) from Allen Mouse Brain Atlas at different stages of mouse brain for *Neurod1*. **h-j**, The genome tracks showing the chromatin accessibility (top), peaks site, peaks coaccessibility (middle), peak-gene linkages, gene tracks (bottom), gene expression (right) for *Neurod2* (h), *Neurod6* (i) and *Igfnpl1* (j) in each cluster. *Neurod1* and *Rfx2* motif were shown as gray box.

Next, we evaluated the MISAR-seq RNA performance during the co-measurement of both modalities. Generally, the total number of UMIs and genes per gird were comparable or slightly better than DBiT-seq in the same 50 μm resolution^12^ (Fig. 1d). The fractions of read-pairs mapped to mitochondrial and ribosome protein of each sample section were lower than those from DBiT-seq (Extended Data Fig. 5a). The tissue permeabilization and fixation is the key bottleneck to ensure the technical compatibility of both modalities. Interestingly, we introduced additional refixation by 4% formaldehyde before mRNA reverse transcription and found this treatment increased the yield and quality of the spatial transcriptome data (Extended Data Fig. 5b and Methods).

**Fig. 5.**
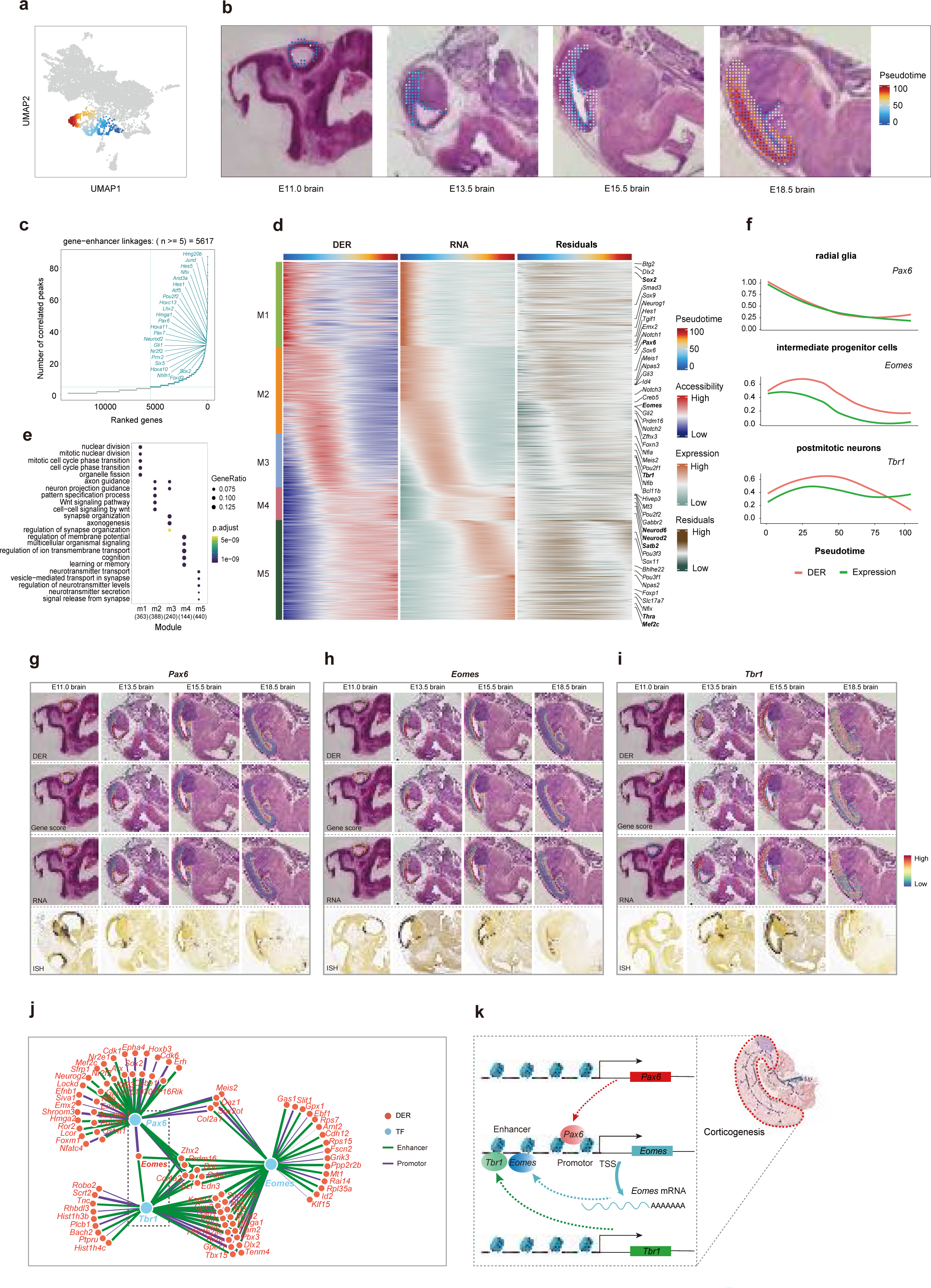
Molecular dynamic and gene regularly network of corticogenesis. **a**, UMAP showing the trajectory of corticogenesis. **b**, Spatial mapping of the trajectory of corticogenesis. **c**, distribution of pseudotime value across development stages. **d**, Scatterplot showing pseudotime value versus distances to the inner layer of cortex, colored by development stages. **e**. The number of significant correlation peaks for each gene (FDR < 0.01 and PCC > 0.1). Some top TF genes are highlighted. **f**, GO enrichment of top 200 highly enhancer-regulated genes. **g**, Heatmap showing domains of enhancer regulatory (DER), gene expression and residual along pseudotime. The residual for each gene was calculated by subtracting min-max normalized gene expression from min-max normalized accessibility activity scores. **h**, GO enrichment of each module gene, top 5 significant GO terms for each module are shown (FDR < 0.01). **I**, Scatterplot of Thra motif enrichment among linked peaks versus correlation between gene expression and accessibility activity score. Highlighted are candidate TF regulators of *Thra* with an absolute regulation score > 1. **j**, Accessibility activity score, and gene expression of *Neurod6*, *Thra*, and *Mef2c* in the pseudotime axis. **k**, Gene regulatory network visualization. Edges are colored by signed regulation score. **l**, Schematic of regulatory relationship among *Neurod6*, *Thra*, and *Mef2c* in the corticogenesis trajectory.

To explore the consistence of the MISAR-seq data on both modalities, the correlation analysis was performed between biological replicates of serial tissue sections for MISAR-seq ATAC data and we found very high data reproducibility (Fig. 1e). Moreover, we also aggregated the chromatin accessibility and mRNA reads from the same E15.5 forebrain (S1) section. Unsurprisingly, the results showed that the RNA reads were located in the 3’ end and the peaks of DNA accessibility were located in the TSS region of the same gene, indicating we could deduce the chromatin-gene expression relationships in cells of the same spatial locations. Besides, the aggregated chromatin profiles from randomly selected 300 spatial grids of an E15.5 forebrain also faithfully reproduced the bulk measurement of chromatin accessibility obtained from the ENCODE data but had better signal-to-noise ratio, suggesting our MISAR-seq approach is robust (Fig. 1f).

The MISAR-seq data was also used to interrogate published scRNA-seq data from the same stage^2^. Take E15.5 brain (S1) as an example, we first performed single cell deconvolution by RCTD to examine the cell type distribution in each spatial grid (see Methods). We found the major cell type distributions have a good agreement with their spatial anatomical annotation (Fig. 1g). We then integrated both MISAR-seq data and scRNA-seq reference data and performed unsupervised clustering (see Methods). 11 clusters were obtained for MISAR-seq ATAC and 14 clusters for MISAR-seq RNA. Expectedly, we found that the cell types agree very well with the brain anatomical region. Then, the clusters were mapped to the spatial section and annotated by spatial anatomic or pseudo-tissue of the reference data separately (Fig. 1h-m). We found that the cell types agree very well with the brain anatomical regions. For example, forebrain dorsal in reference single-cell dataset were mapped to regions of the forebrain dorsal pallium intermediate, ventricular and superficial (Fig. 1j,m).

### Profiling of chromatin accessibility and gene expression in development embryo mouse brain with MISAR-seq

Having the co-assay of spatial ATAC and RNA at E11.0, E13.5, E15.5 and E18.5 stages, we sought to examine the comprehensive view of both parameters along the brain developmental process. After filtering out non-tissue grids, a total of 7118 grids with both chromatin accessibility and gene expression were obtained. To accommodate the integrative analysis of both chromatin accessibility and transcriptome, we aligned cross-data in parallel using slices of MISAR-seq ATAC. First, for ATAC profiles, a pixel by tile matrix was generated by aggregating reads in 5 kilobase bins across the mouse genome and an ATAC ArchR object was created. For RNA profiles, we use addGeneExpressionMatrix function to add RNA profiles in the ArchR object. Then, latent semantic indexing (LSI) was applied for dimensionality reduction and Harmony was used to remove technical batch effects (see Methods). Setting a basic layer of comparison, a manual anatomical annotation was performed to reveal major tissue organization following Kaufman’s Atlas of Mouse Development and Allen Brain Atlas (Fig. 2h). We found both spatial-ATAC clusters and spatial-RNA-seq clusters can faithfully recapitulate the tissue organization in the developing mouse brain, with slightly different in minor details (Fig. 2). For example, among the 11 ATAC or RNA clusters, ATAC cluster 2 or RNA cluster 4 represents cartilage in the mouse embryo. ATAC cluster 5 and RNA cluster 5 are associated with forebrain of dorsal pallium (DPallm), including the ventricular and mantle region of dorsal pallium. At E11.0, the RNA profile showed higher resolution in defining the spatial structures compared to ATAC information. Overall, the spatial ATAC and RNA clusters correlated well, which was further demonstrated in the joint spatial clustering with integrated datasets. The combined clusters also showed that the spatial complexity was increased gradually along the brain development, with more discrete cell clusters emerged at E18.5 (Fig. 2e-g). Besides, the analysis from replicate sections obtained similar results (Extended Data Fig. 6). Together, our data shows that major anatomical structures can be readily delineated by spatially resolved chromatin and transcriptome profiling.

### Integrated analysis of chromatin accessibility and gene expression across brain tissue types

To fully reveal the relationship and molecular mechanisms between chromatin state and gene expression, the chromatin- and transcriptome-based spatial cell clustering were compared. We first evaluated the consistency between the cell clusters based on independent modality and found a high fraction of cells were shared by respective ATAC and RNA clusters (Fig. 3a). By connecting the ATAC clusters with RNA clusters, we found that the overall patterns of chromatin accessibility were associated with more diverse gene expression clusters (Fig. 3a,b), indicating a lower clustering power for ATAC information possibly owing to the sparsity of chromatin data that are still undergoing maturation. For example, two ATAC clusters 2 and 9 which are mostly present at E11.0 were evolved into many RNA clusters at late stages. We also found that the grid composition of the clusters became more distinct during brain development, indicating an increased consolidation of tissue identity (Fig. 3c).

Next, we aimed to find the signature features related to each cluster. With ArchR, we are able to generate “gene activity score” to enable accurate cluster annotation (see Methods). In summary, we identified 571 marker genes in ATAC-based clusters and 2473 marker genes in RNA-based clusters across the integrated 14 tissue types, in which, many of these markers showed cluster-specific expression and can be validated by in situ hybridization data from Allen Mouse Brain Atlas (Fig. 3d,e and Extended Data Fig. 7). We selected the top15 marker genes to perform Gene Ontology (GO) analysis, and found the annotations were consistent between two modalities and aligned well with their anatomical structures. Furthermore, as reported in other dual-omics assay, the cell-type identities of the major clusters were better supported by specific accessible chromatin accessibility in promoter regions of marker genes (Fig. 3f)^15^.

### MISAR-seq identified regulatory machinery for mouse brain development

To disentangle the complexity of gene regulatory networks that drive brain development, we sought to identify the regulatory mechanisms that function for establishing the spatial organization of the mouse brain. To this end, we first created pseudo-bulk replicates based on the combined clusters using the addReproduciblePeakSet function of ArchR and called peaks with MACS2 considering per-cell ATAC-seq data is essentially binary (accessible or not accessible) (see Methods). We counted the number of peaks and found that there are high proportion of promoter and distal peaks in each cluster, indicating that a lot of regulatory elements were obtained in our MISAR-seq ATAC data (Extended Data Fig. 8a). We identified a total of 30272 signature peaks enriched in each cluster and labeled the top 4 peaks in the heatmap, in which, most clusters had high numbers of differential peaks (Extended Data Fig. 8b).

Next, we performed motif enrichment analysis based on differential peaks of each cluster with motif database CISBP(v2) using the function of peakAnnoEnrichment from ArchR (see Methods). We obtained 37 enriched motifs in total and found the corresponded transcription factors (TF) showed a tissue-specific distribution (Fig. 4a). For example, motifs of *Myod1*, *Myog* and *Myf5* were enriched in cluster 1 where represents the muscle cells^16^. Motifs of *Neurod1*, *Neurod4* and *Neurod6* were highly enriched in cluster 8 of DPallm, corroborating their reported role in regulating neuronal progenitor differentiation and neuronal specification in the cerebral cortex^17^. Interestingly, the cluster 8 also possess the highest abundance of peaks and motifs, suggesting a predominant presence in mice brain development (Fig. 4a and Extended Data Fig. 8b).

To further predict the TF enrichment on a per-grid basis from the sparse chromatin accessibility data, we calculated the variability deviation for the enriched motifs (see Methods). Interestingly, after ranked the TFs by variable deviation, we found the top 21 TFs could be roughly classified into a few major TF families, including *Rfx* family, *Runx* family and basic Helix-loop-Helix (bHLH) family. The bHLH family members such as *Ascl1*, *Ascl2*, *Neurod1*, *Neurod4*, *Neurod6* and *Atoh1* were well-known master regulators of cell proliferation and neuronal specification^16^. *Rfx* TFs were reported to have both redundant and specific functions in ependymal cell of the brain ventricle^18^. To further uncover the detailed regulatory mechanisms under mouse brain development, we performed peak-gene lineage analysis and identified 11772 peak-gene linkages by ArchR and these linkages were divided into 15 modules according to the pattern of chromatin accessibility and gene expression (Fig. 4c and Methods). Not surprisingly, the chromatin accessibility modules and transcriptome modules have a good consistence with each other. To reveal the potential regulatory function of those peak-gene linkages in brain, the top ranked TFs of *Neurod1* and *Rfx2* were selected to represent the regulation in cortex region or in the diencephalon and hindbrain region, respectively (Fig. 4a,b). The TF footprinting analysis of *Neurod1* and *Rfx2* predicted their enriched binding locations for the corresponding brain regions (Fig. 4d and Extended Data Fig.10b). The region-specific promoter activity of these TFs can be verified in the chromatin accessibility, gene expression, TF deviation score and in situ hybridization data from Allen Mouse Brain (Fig. 4e-g and Extended Data Fig. 10a). After scanning the motif of Neurod1 and Rfx2 in all peak-gene linkages, we obtained 328 motif sites for *Neurod1* and 697 motif sites for *Rfx2*. We showed the significant motif sites for *Neurod1* and *Rfx2* (peak-gene correlation >= 0.7, Fig. 4b,c). Unexpectedly, we found *Neurod2*, *Dync1i1*, *Neurod6* and *Igfbpl1* were enriched in DPall regions for *Neurod1* motif and *Igfbpl1*, *Dpysl5*, *Nyap1*, *Atp1a3* and *Pou2f2* were enriched in DPall, diencephalon and hindbrain for *Rfx2* motif (Fig. 4c), indicating the TFs function have a high correlation with their tissue location. The genome track plot showed that *Neurod1* motif was located at the promoter or enhancer region of *Neurod2* and *Neurod6*, suggesting both of them can be directly regulated by *Neurod1*^19^. Meanwhile, We also identified other dynamically regulated promoter interactions with specific enhancers by peak gene linkages and peak co-accessibility analysis (Fig. 4h,i and Extended Data Fig. 9e). Besides, the Neurod1-regulated target genes also showed region-specific expression in DPall regions (Extended Data Fig. 7e,10d and 10e). These are also reflected in the *Rfx2 motif and Rfx2*-regulated targets e.g. *Pou2f2*, *Nyap1*,

*Atp1a3* and *Dpysl5* (Extended Data Fig. 9a-d, Extended Data Fig. 11). Interestingly, we found *Igfbl1*, the member of insulin-like growth factor binding proteins, can be regulated by both *Neurod1* and *Rfx2* because both motifs were found in its promoter or enhancer region. The chromatin accessibility and gene expression of *Igfbpl1* proved the specific activation in DPall region (Fig. 4j and Extended Data Fig. 10c), in line with the reported role of *Igfbpl1* as important regulator in the development of mouse forebrain^20^.

All together, we uncovered the signature TFs for each cluster based on the differential peaks and identified key regulators that enrich in specific brain regions using peak-gene expression correlation revealed by the co-mapping of spatial ATAC and spatial RNA profile.

### Molecular dynamic and gene regulatory network of corticogenesis

Corticogenesis is a complex development process characterized by the proliferation, differentiation, and migration of cortex progenitor cells, and the inside-out gradient of neurogenesis. The regulatory mechanism of cis-regulatory elements such as enhancers and promoters are central for cortical neuron cell fate determination and laminar structure organization. Empowered by the unique ability of joint mapping the spatial ATAC and RNA data, we anticipated that asynchronous cell fate determination and cascade regulation may be illustrated through the integration of both modalities in a spatial and temporal manner.

We first selected the combined peak accessibility and gene expression that map to the cortex regions from the united UMAP clusters to infer the corticogenesis trajectory (Fig. 5a, b, see Methods). Of note, the pseudotime axis captured well the spatiotemporal changes in cortex development (Fig. 5c, d and Methods). To connect the enhancer elements and gene expression, we applied a correlation-based method to the smooth pseudotime axis of the combined manifold (see Methods). We identified 66,805 putative enhancer-to-gene linkages. Interestingly, highly enhancer-regulated genes (genes with a large number of peaks-gene linkages) (blue color in Fig. 5e) are associated with forebrain development and neuron differentiation. For example, transcription factors such as *Sox2*, *Dlx2*, and *Hes* showed high expression in early phase of the pseudotime (Fig. 5g). *Sox2* has a critical role in maintaining neuron stem cells^21^. *Hes* genes are downstream targets of the *Notch* signaling pathway that represses proneural genes and maintain neuron stem cells boundaries^22^. *Dlx* genes can function as modulators to determine neuron versus oligodendrocyte cell fate^23^. Therefore, these results indicate that the spatiotemporal trajectory analysis has the potential to identify important regulatory mechanisms for early cortex cell fate determination.

To further analyze the dynamics of accessibility activity and gene expression along the corticogenesis trajectory, we computed the domain of enhancer regulatory (DER) activity by aggregating linked peak counts for each gene (see Methods). The generalized linear model was fitted on each gene expression and accessibility activity score to select the pseudotime-dependent features. We identified 1160 dynamic regulatory interaction pairs and clustered them into 5 modules (Fig. 5g). Modules along the pseudotime were enriched in functions such as cell proliferation, neuron migration, axonogenesis, neuron projection, and synapse activity just in the developmental order of corticogenesis (Fig. 5h). Coordination of these interaction pairs indicates that a set of highly regulatory genes drive cell fate decision and determine lineages differentiation during corticogenesis. Of note, we observed a predominately asynchronous pattern between peak accessibility and gene expression at late pseudotime. For example, we noticed module 5 overall gained accessibility earlier than their gene expression apparently.

Next, we utilized FigR to infer the transcriptional regulators of target genes^24^. Integrated peak-to-gene links and motif enrichment in linked peak regions allow us to define many putative activators and repressors and construct the gene regulation network (Fig. 5i-l and Extended Data Fig. 12a). We found thyroid hormone receptor *Thra* is a node TF that connects many targets. Previously, thyroid hormone receptors have been suggested to play a crucial role in human and mouse cortical neurogenesis^25, 26^. However, the regulatory mechanism remains unknown. By integrating peak accessibility and gene expression, we found that motifs of *Neurod6* and *Mef2c* were enrichment in linked peaks of *Thra*, suggesting *Thra* may be positively regulated by *Neurod6* and *Mef2c* (Fig. 5i). Ordering along the pseudotime, *Neurod6* accessibility was increased first, followed by Thra accessibility, *Neurod6* RNA, and *Thra* RNA. Finally, *Mef2c* accessibility and RNA were increased toward the pseudotime endpoint (Fig. 5j). Interestingly, we also found that *Neurod6*, *Thra*, and *Mef2c* TFs show high centrality in the TF-targets network and are positively correlated with each other (Fig. 5k), pointing to a potential TF regulatory cascade during cortex development^27^(Extended Data Fig. 12b). Hence, MISAR-seq has the power to map the gene regulatory logic of mouse corticogenesis and provides new insights for decoding the spatiotemporal dynamic of tissue organization.

## Discussion

Recently, spatially resolved transcriptomics has been a technological frontier that reshapes biomedical research with emergence of many new methodologies and applications. Importantly, the architecture and tissue function are regulated across multiple layers, including epigenetic regulations such as modulation of gene expression by changes in histone modification and chromatin accessibility. Therefore, it is highly desirable to systematically examine the spatial gene expression together with epigenetic information directly in the same cell populations. In the current study, we reported a high-throughput method for simultaneous profiling of chromatin accessibility and gene expression with spatial information retained. We demonstrated the utility of MISAR-seq by applying it to the mouse embryonic brain development at four developmental stages. Our results show that MISAR-seq can identify the chromatin potential and gene expression of the same spatial location. The integrative analysis enables us to define the cell fate determination of mouse fetal brain, and uncover the spatiotemporal regulatory principles during mammalian brain development.

It worth noting that currently the method has not achieved single-cell resolution, therefore making the defined interpretation of regulatory mechanisms or trajectory inference still challenging, considering pronounced heterogeneity exists in the spatial organization of diverse biological systems. Although we only presented data obtained on embryonic brain tissues, we have adapted the method to other tissue types such as tumor samples and achieved similar performance. Considering similar principle has been implemented in a recent spatial-ATAC-seq method and robust results can be obtained^10^, we anticipate the MISAR-seq would be compatible for a wide range of tissue type. Future optimization to further increase the spatial resolution, the scalability and additional layers of omics information will provide versatile tools to disentangle the complexity of gene regulatory and cell–cell communication networks that drive cell functions and responses in structurally organized tissues.

## Author contributions

G.P., and J.F. designed the study. J.F., X. Z., Y.Q., L.W. and S.Q. performed the experiments.

J.F. and M.Z. analyzed the data with help from F.Q., G.C. and Z.L. K.C. provided reagents and suggestions. J.F. and G.P. wrote the manuscript with the help of others.

## Acknowledgments

This work was supported in part by National Key R&D Program of China (2018YFA0107200, 2018YFA0801402), the "Strategic Priority Research Program" of the Chinese Academy of Sciences (XDA16020404), National Natural Science Foundation of China (31871456 and 32100483), Guangdong Basic and Applied Basic Research Foundation (2019B151502054, 2019A1515110985 and 2020A1515110517), Frontier Research Program of Bioland Laboratory (Guangzhou Regenerative Medicine and Health Guangdong Laboratory, 2018GZR110105013), Jiazi Research Innovative Project of Bioland Laboratory (2019GZR110108001), Science and Technology Planning Project of Guangdong Province (2020B1212060052). We thank Patrick Tam for discussions and critical reading of this study.

---

**Extended Data Fig. 1.**
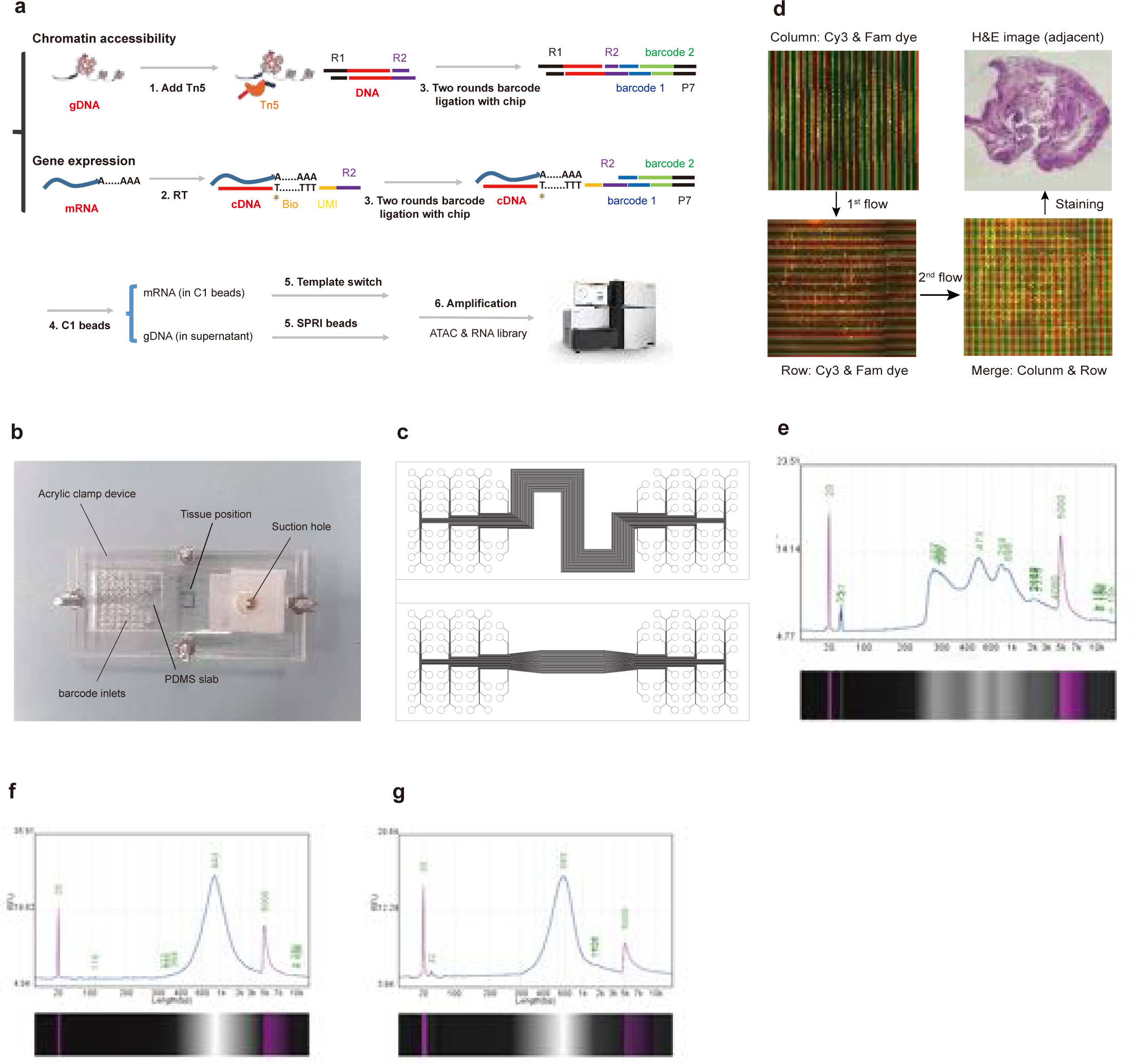
MISAR-seq: workflow, design and library distribution. **a**, MISAR-seq workflow. The tissue section was first subjected to Tn5 fragmentation, followed by reverse transcription (RT) and finally two rounds of barcode (50 kinds each) ligation on chip. R1, Read1 adaptor. R2, Read2 adaptor. **b**, The composition of microfluidic devices used in this article, including PDMS slab, barcode inlets, acrylic clamp and suction device. **c**, AutoCAD design of PDMS chip with 50μm channel width. **d**, Verification of leakage or diffusion by Cy3 and Fam dye. **e-g**, Size distributions of ATAC library (e), cDNA amplicons (f) and cDNA library (g).

**Extended Data Fig. 2.**
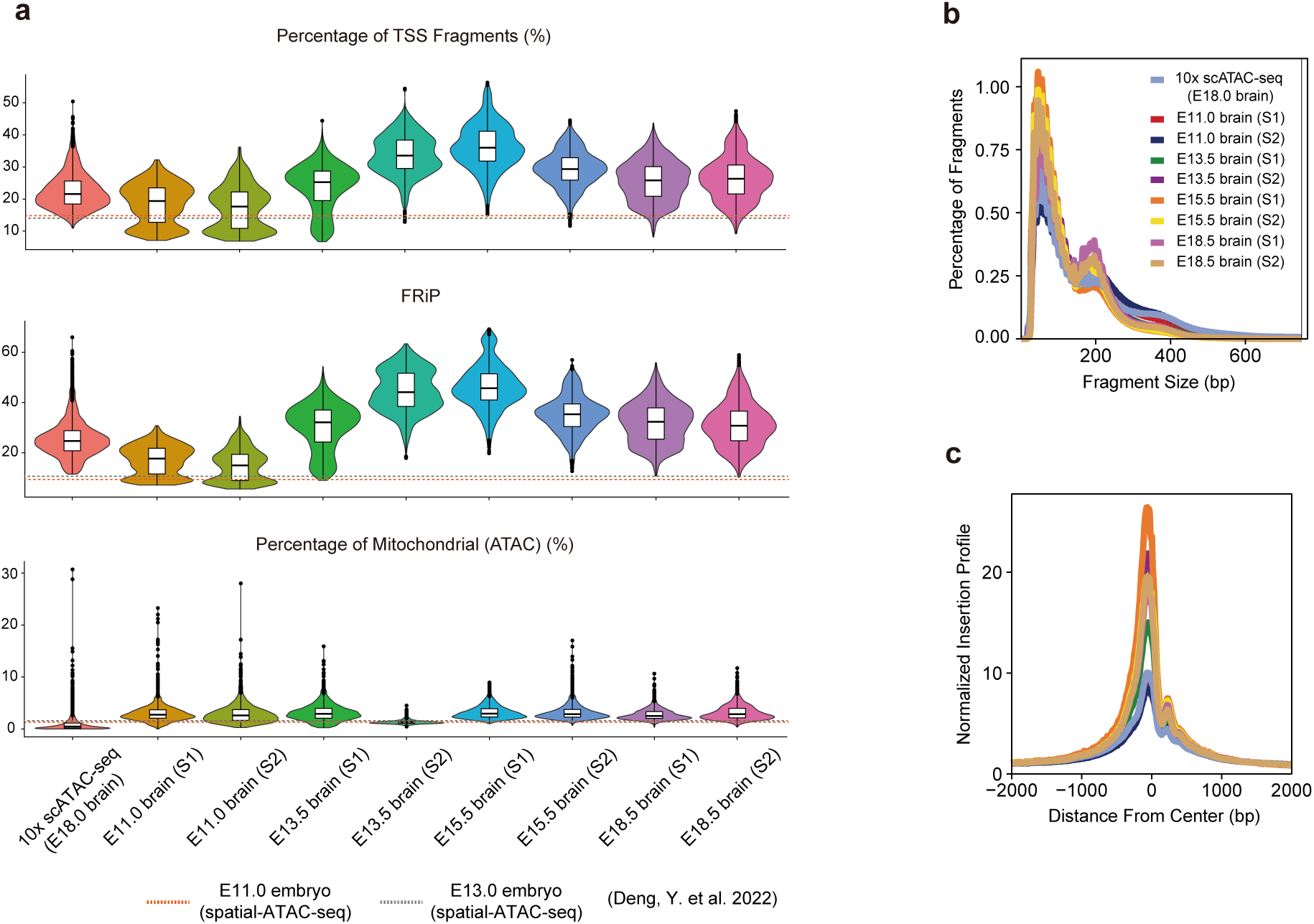
Data quality of spatial ATAC in MISAR-seq. **a**, Comparison of percentage of mitochondrial, TSS fragments and FRiP value between MISAR-seq and 10x scATAC-seq, S1, Section1. S2, Section2. Dashed lines indicate the results from Spatial-ATAC-seq^10^.. **b**-**c**, Comparison of insert fragments size distribution (b) and TSS enrichment profiles (c) between MISAR-seq and 10x scATAC-seq.

**Extended Data Fig. 3.**
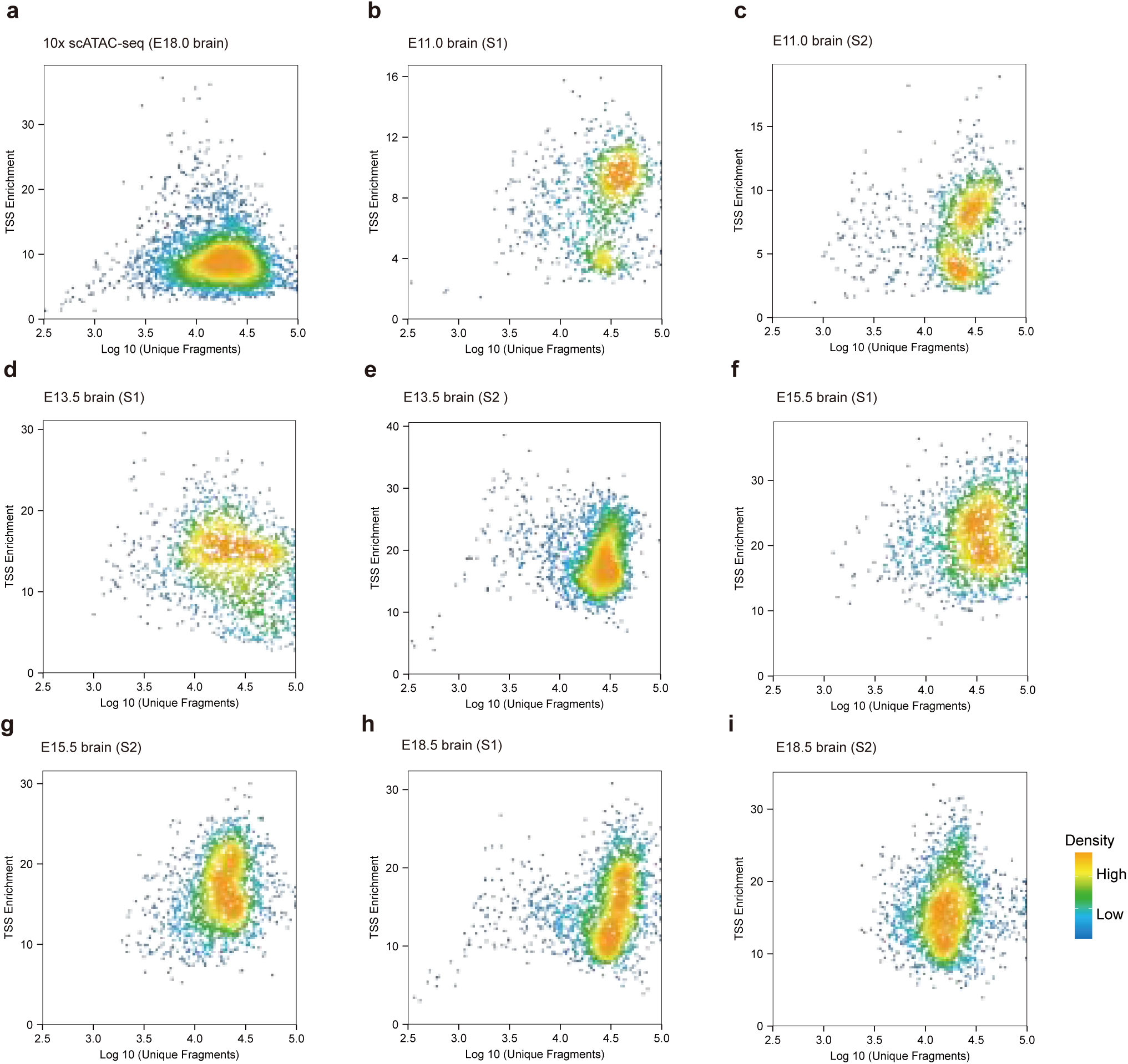
TSS enrichment in MISAR-seq. **a-i**, Comparison of TSS enrichment scatterplot between 10x scATAC-seq (a) and different stages of mouse brain from MISAR-seq (b-i).

**Extended Data Fig. 4.**
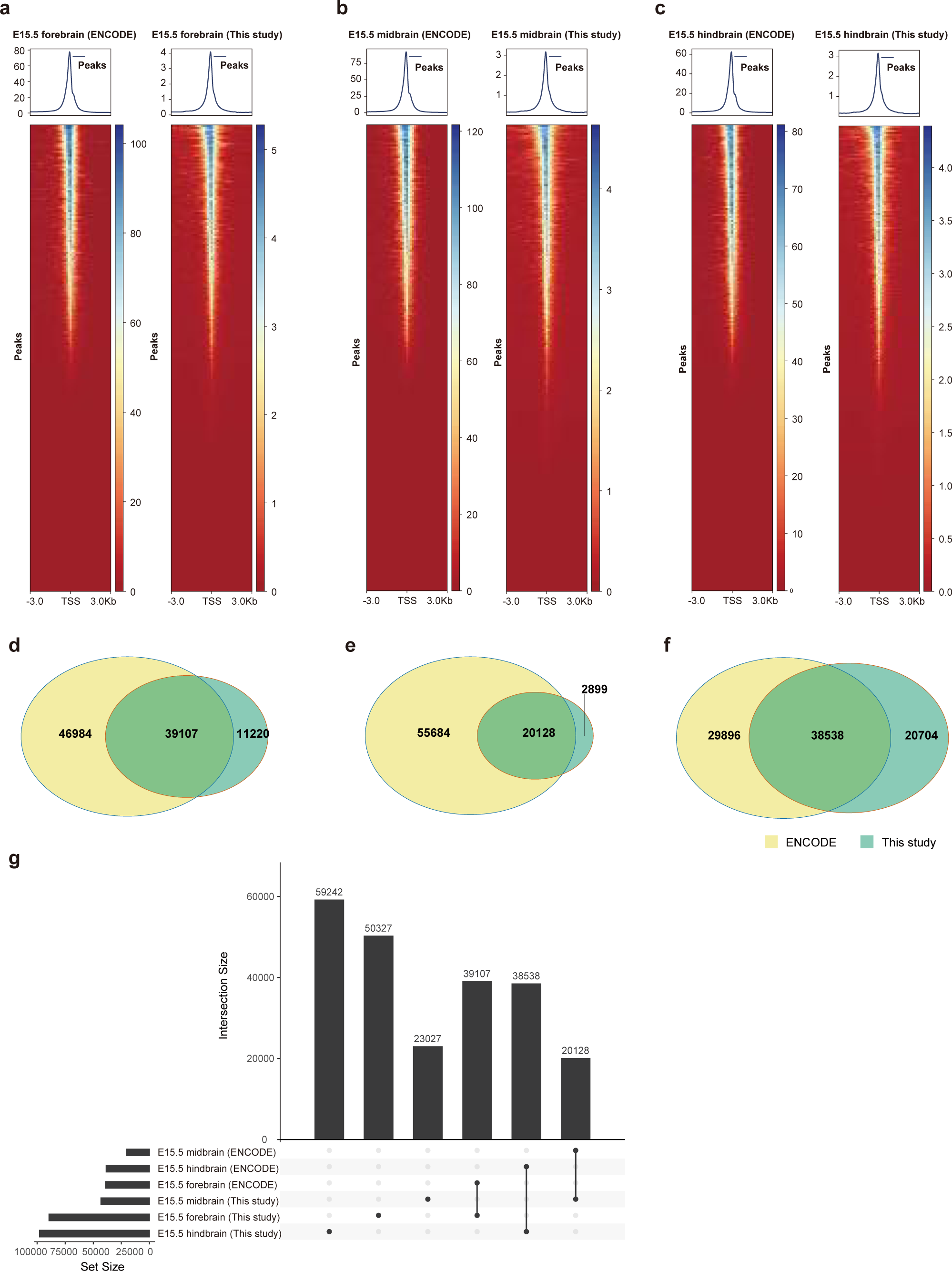
Comparison of peaks called with MISAR-seq and ENCODE bulk ATAC E15.5 brain data. **a-c**, Comparison of aggregated forebrain (a), midbrain (b) and hind-brain regions (c) of peak-centered heatmap from MISAR-seq and bulk corresponding brain region from ENCODE bulk ATAC E15.5 brain data. **d-f**, The overlap of peaks called from aggregated forebrain (d), midbrain (e) and hindbrain regions (f) of Venn diagram from MISAR-seq and ENCODE datasets. **g**, Quantitative analysis of peaks upset plot called from aggregated forebrain, midbrain and hindbrain regions from MISAR-seq and ENCODE datasets.

**Extended Data Fig. 5.**
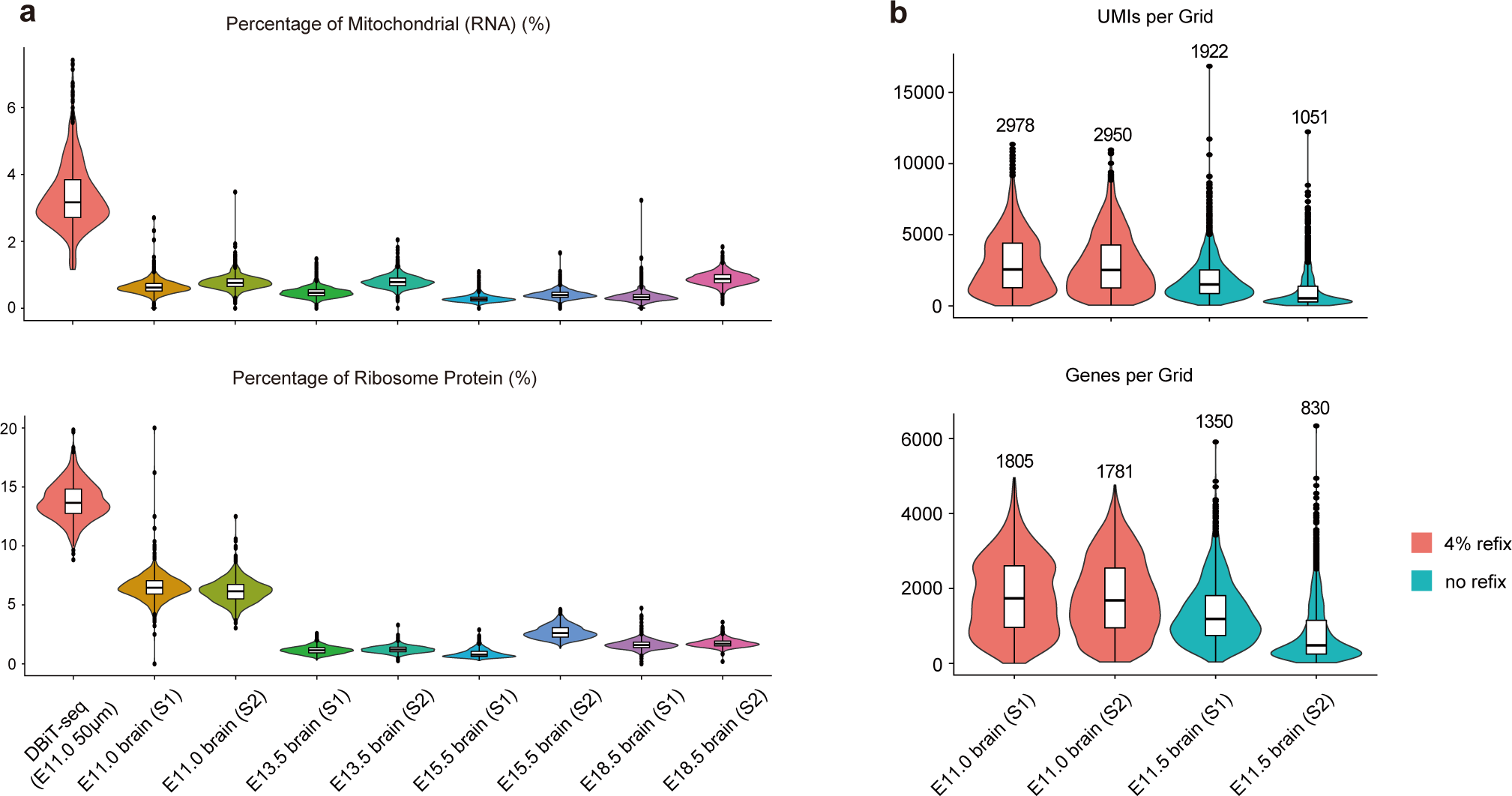
Data quality of spatial RNA in MISAR-seq. **a**, Comparison of percentage of mitochondrial and ribosome protein per grid between MISAR-seq and DBiT-seq. **b**, Comparison of UMIs and genes number per gird between whether or not add 4% formaldehyde after Tn5 fragmentation in MISAR-seq workflow.

**Extended Data Fig. 6.**
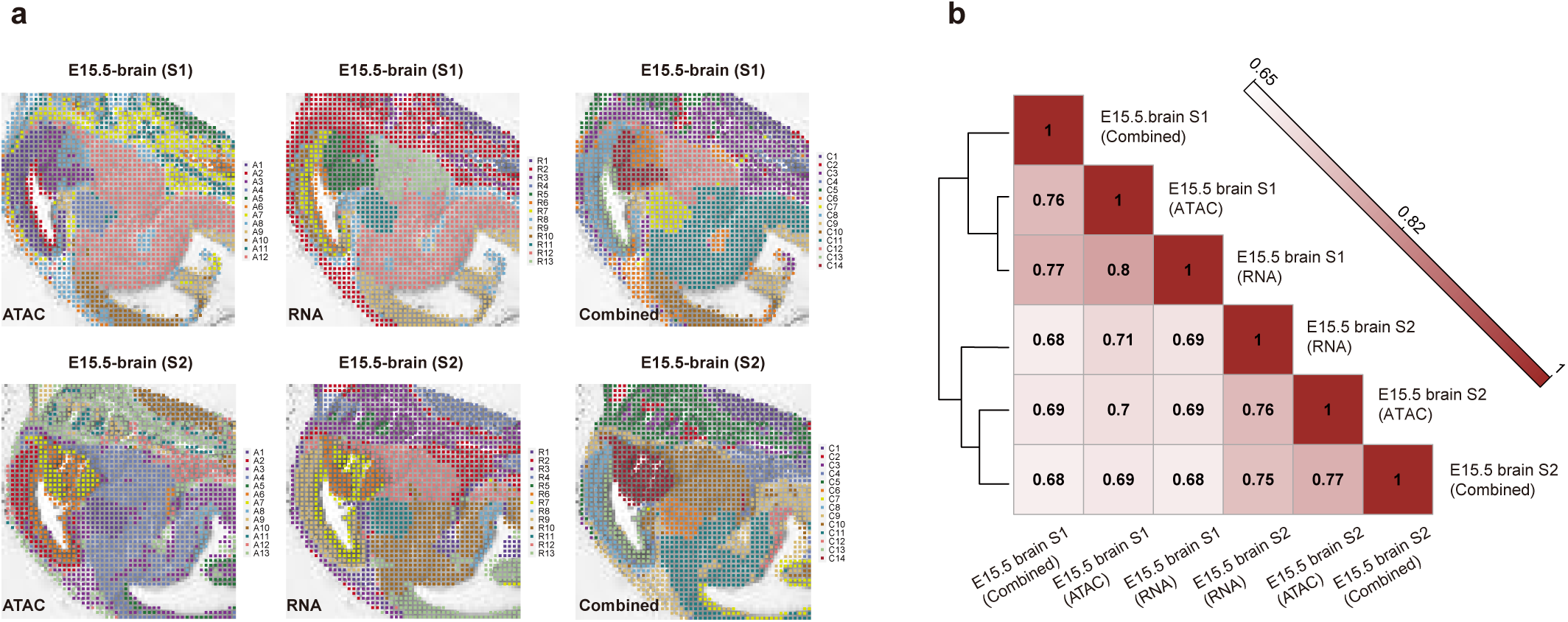
Spatial chromatin accessibility, gene expression and combined mapping for mouse brain development at E11.0, E13.5, E15.5 and E18.5 from section2. **a-c**, Spatial ATAC (a), RNA (b) and combined (c) UMAP visualization of different mouse brain development stage, colored by different clusters. **d**, Combined spatial ATAC and RNA UMAP visualization of integrated different mouse brain development stage, colored by different sample sections. **e-g**, Unsupervised clustering of mouse brain sections for spatial ATAC (e), RNA (f) and combined (g).

**Extended Data Fig. 7.**
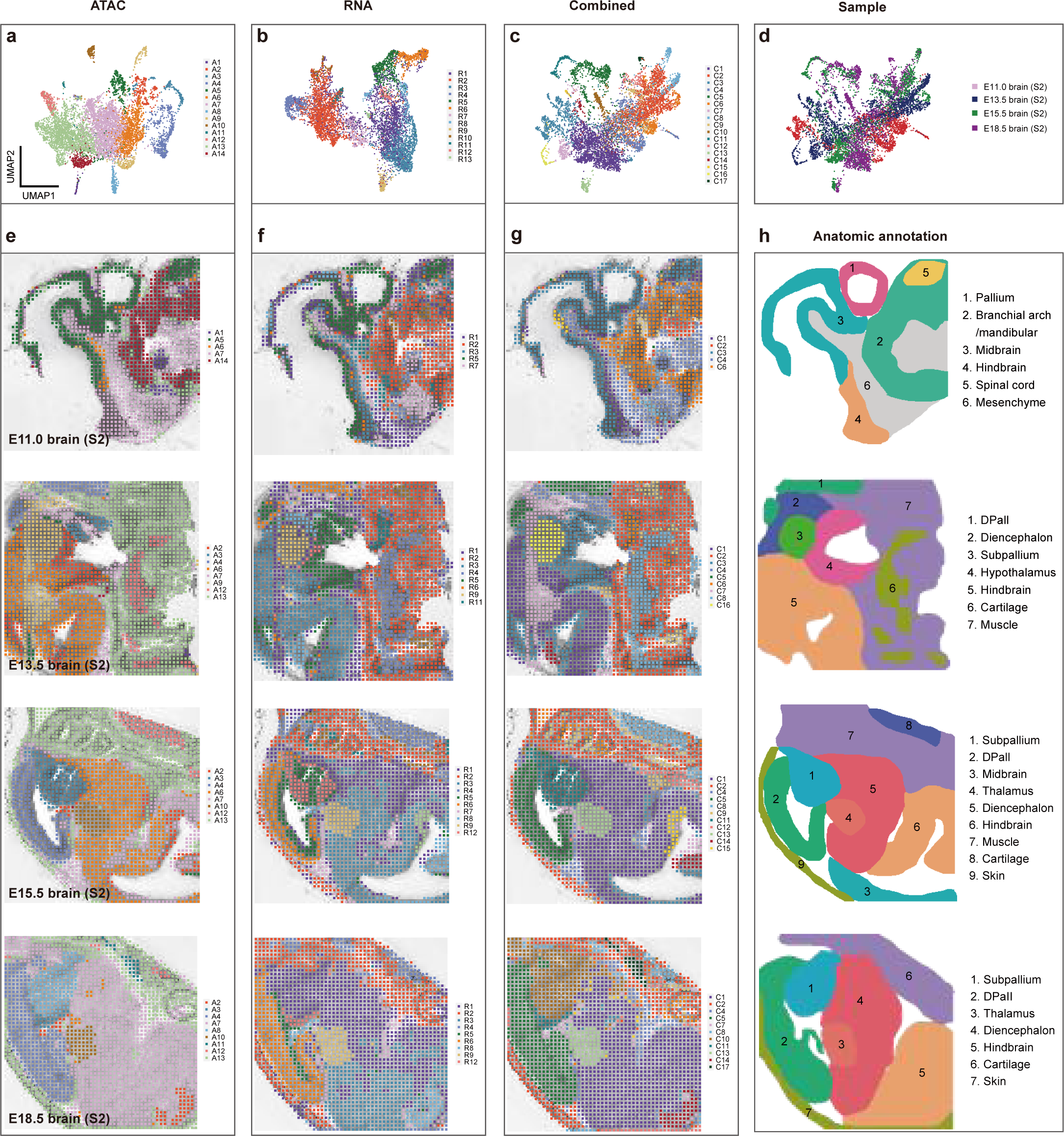
Comparison of spatial expression of selected genes with in situ hybridization data from Allen Mouse Brain Atlas. **a-f**, Spatial mapping of gene expression (RNA) and gene score (ATAC) for selected marker genes, and in situ hybridization of corresponding genes at E11.0, E13.5, E15.5 and E18.5 mouse brain.

**Extended Data Fig. 8.**
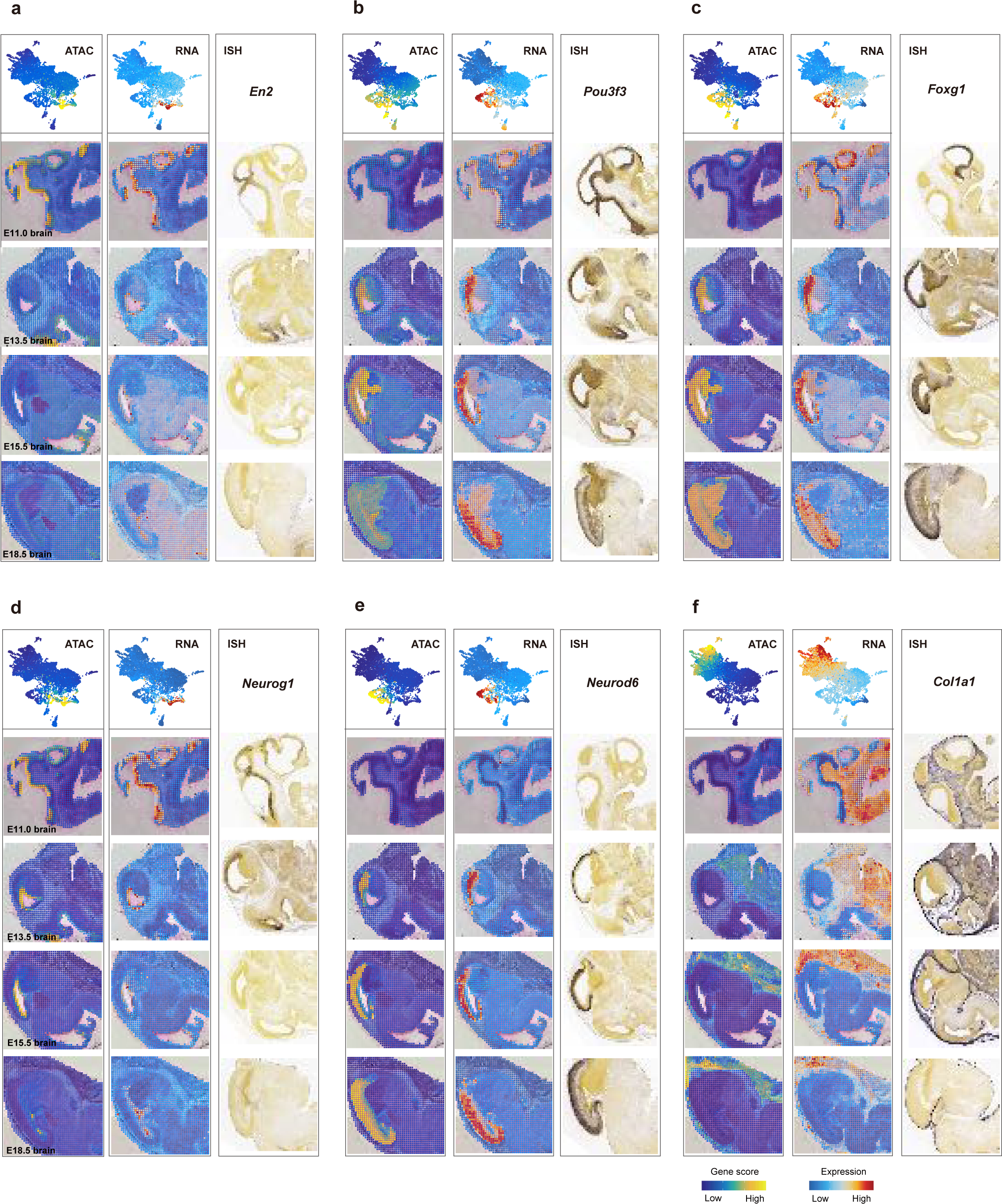
Marker genes analysis for each cluster. **a**, Peak annotation and proportion plot for each cluster. **b**, Heatmap of spatial ATAC marker peaks across all clusters.

**Extended Data Fig. 9.**
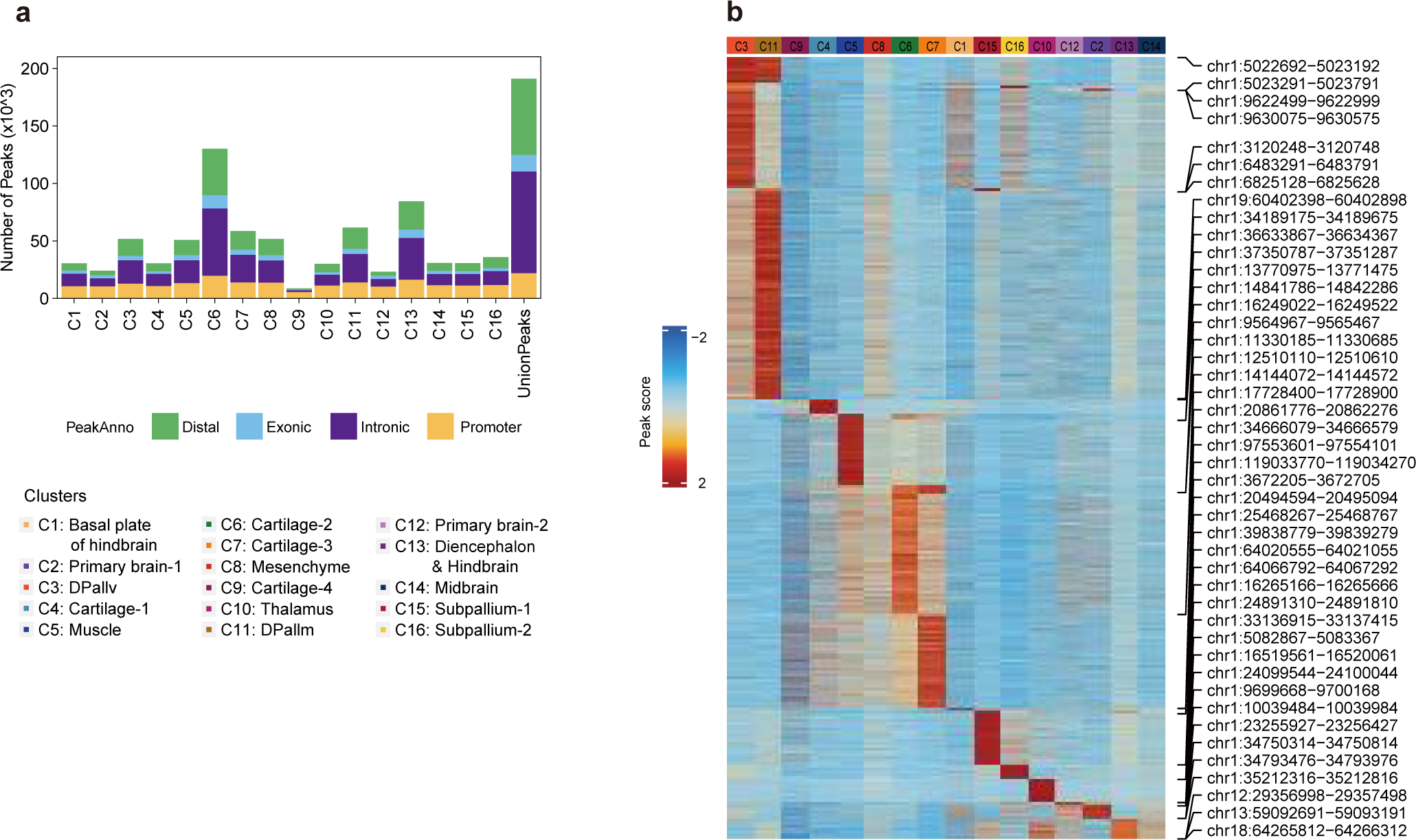
The genome tracks of representative target genes. **a-e**, The genome tracks showing the chromatin accessibility (top), peaks site, peaks coaccessibility (middle), peak-gene linkages, gene tracks (bottom), gene expression (right) for *Pou2f2* (a), *Nyap1* (b), *Atp1a3* (c), *Dpysl5* (d) and *Dync1i1* (e) in each cluster. *Neurod1* and *Rfx2* motif were shown as gray box.

**Extended Data Fig. 10.**
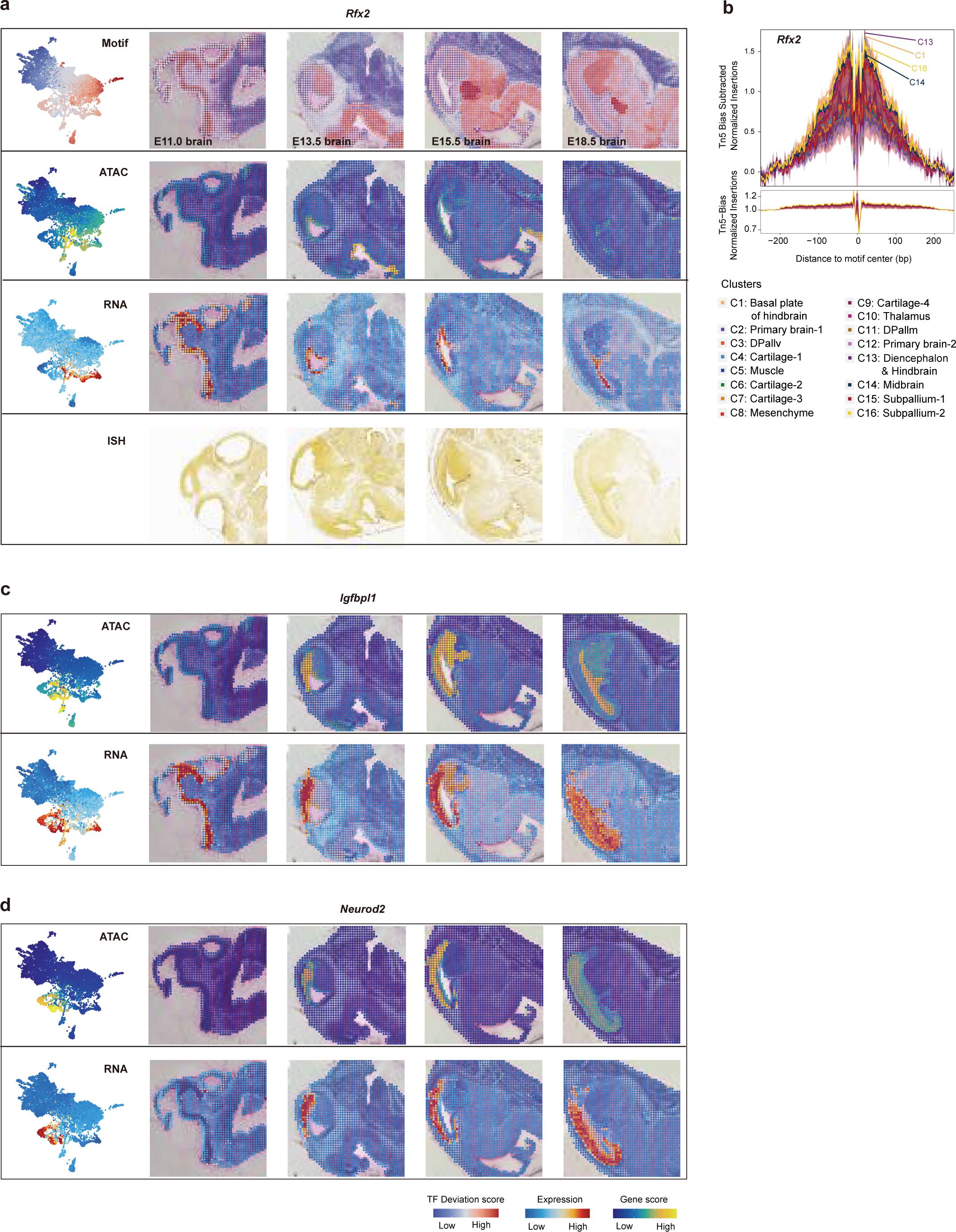
The UMAP embedding and spatial mapping of gene scores, gene expression and deviation scores for represented gene. **a**, UMAP embedding and spatial mapping of gene scores, gene expression, TF deviation score and in situ hybridization results from Allen Mouse Brain Atlas at different stages of mouse brain for *Rfx2*. **b**, Tn5 bias-adjusted transcription factor footprints for *Neurod1* motifs. **c-e**, UMAP embedding and spatial mapping of gene scores and gene expression for *Igfbpl1* (c), *Neurod2* (d) and *Dync1i1* (e).

**Extended Data Fig. 11.**
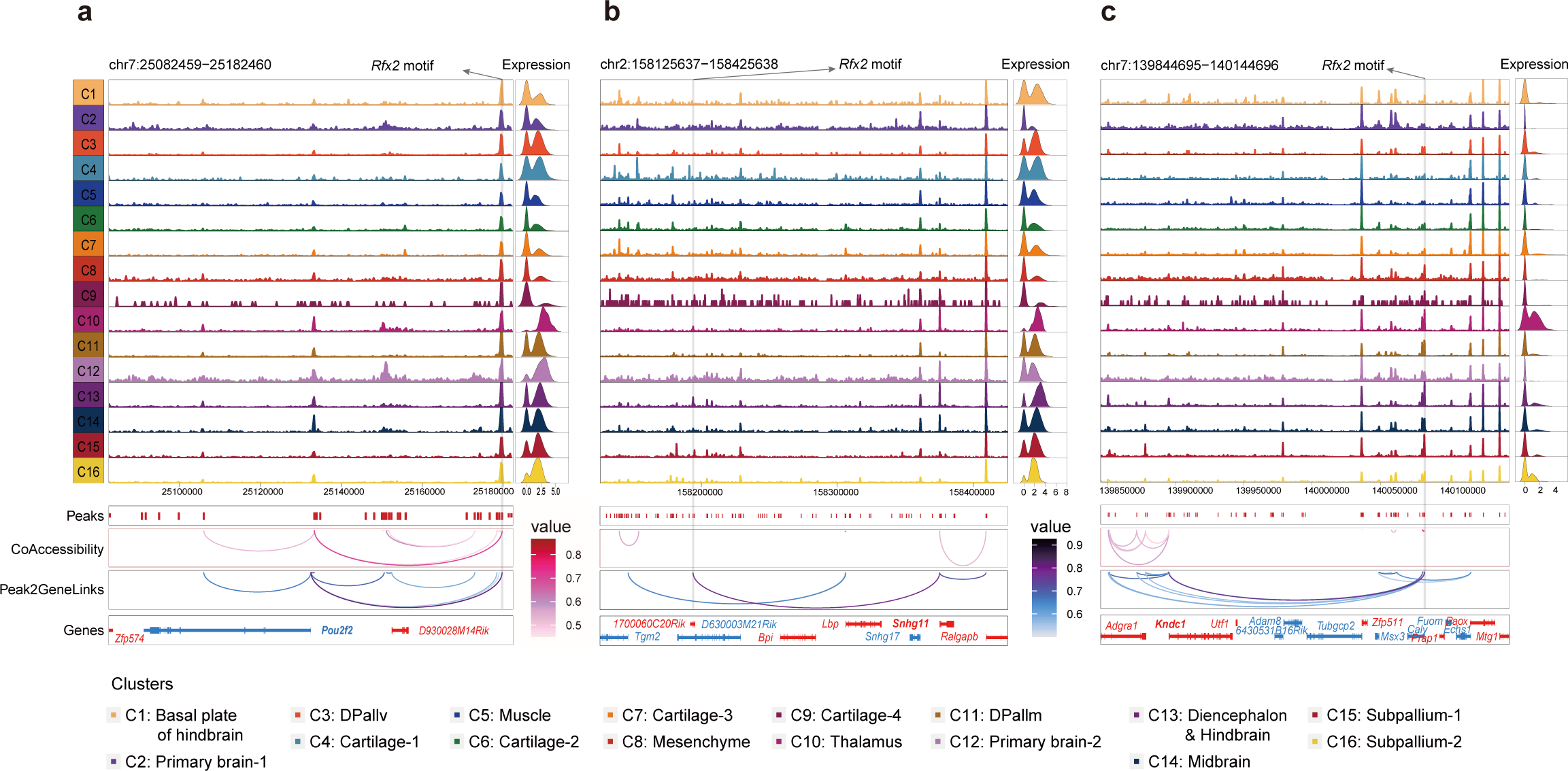
The UMAP embedding and spatial mapping of gene scores, gene expression and deviation scores for represented gene. **a**, UMAP embedding and spatial mapping of gene scores, gene expression, TF deviation score and in situ hybridization results from Allen Mouse Brain Atlas at different stages of mouse brain for *Pou2f2*. **b-d**, UMAP embedding and spatial mapping of gene scores and gene expression for *Nyap1* (b), *Dpysl5* (d) and *Atp1a3* (d).

**Extended Data Fig. 12.**
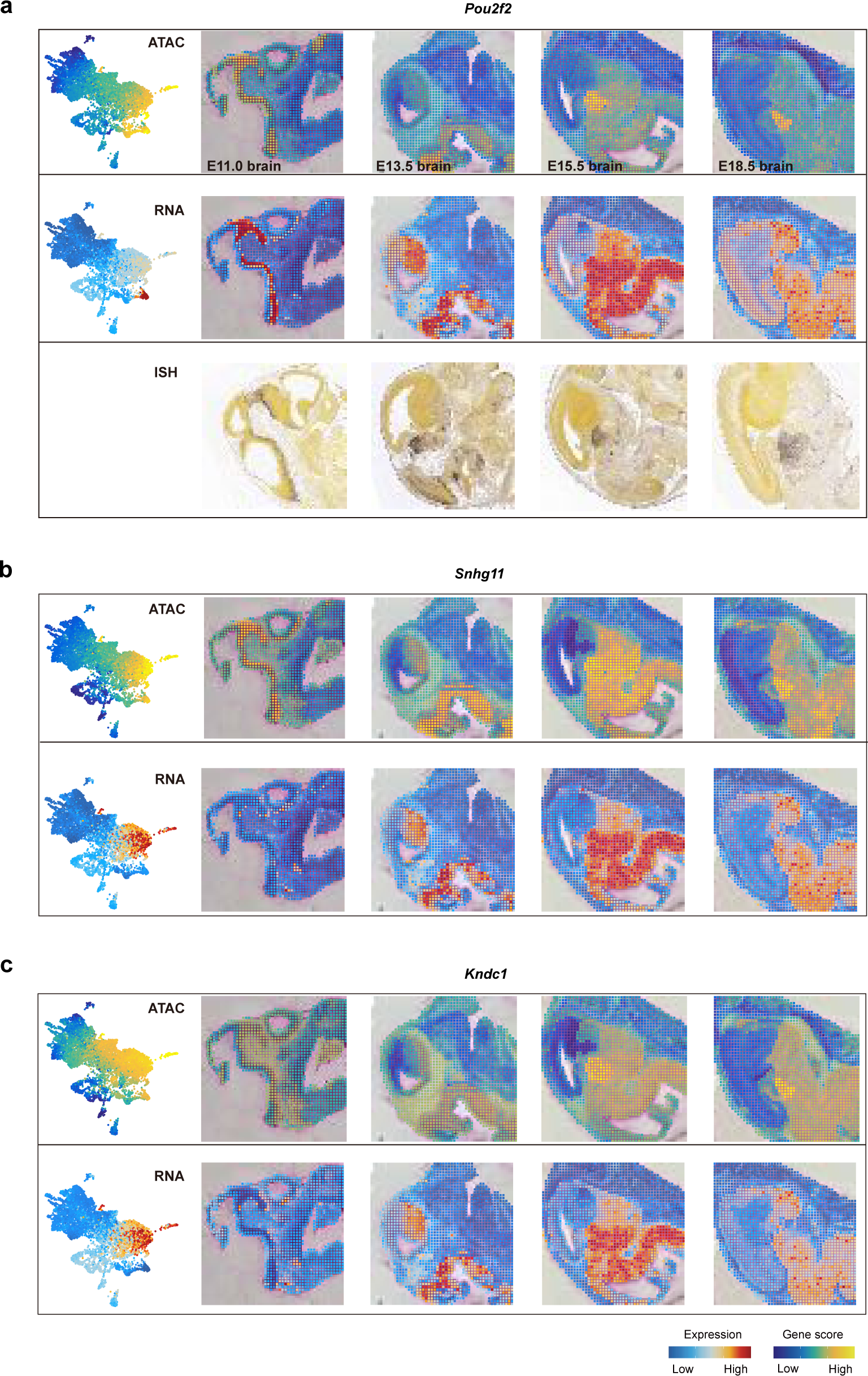
Putative activators and repressors in cortical development and in situ hybridization images of *Thra* and *Mef2c*. **a**, TFs ranked by mean regulation score, highlighting top20 putative TF activators (right) versus TF repressors (left). **b-c**, In situ hybridization and expression images of *Thra* (b) and *Mef2c* (c) downloaded from Allen Mouse Brain Atlas.

**Extended Data Fig. 13.**
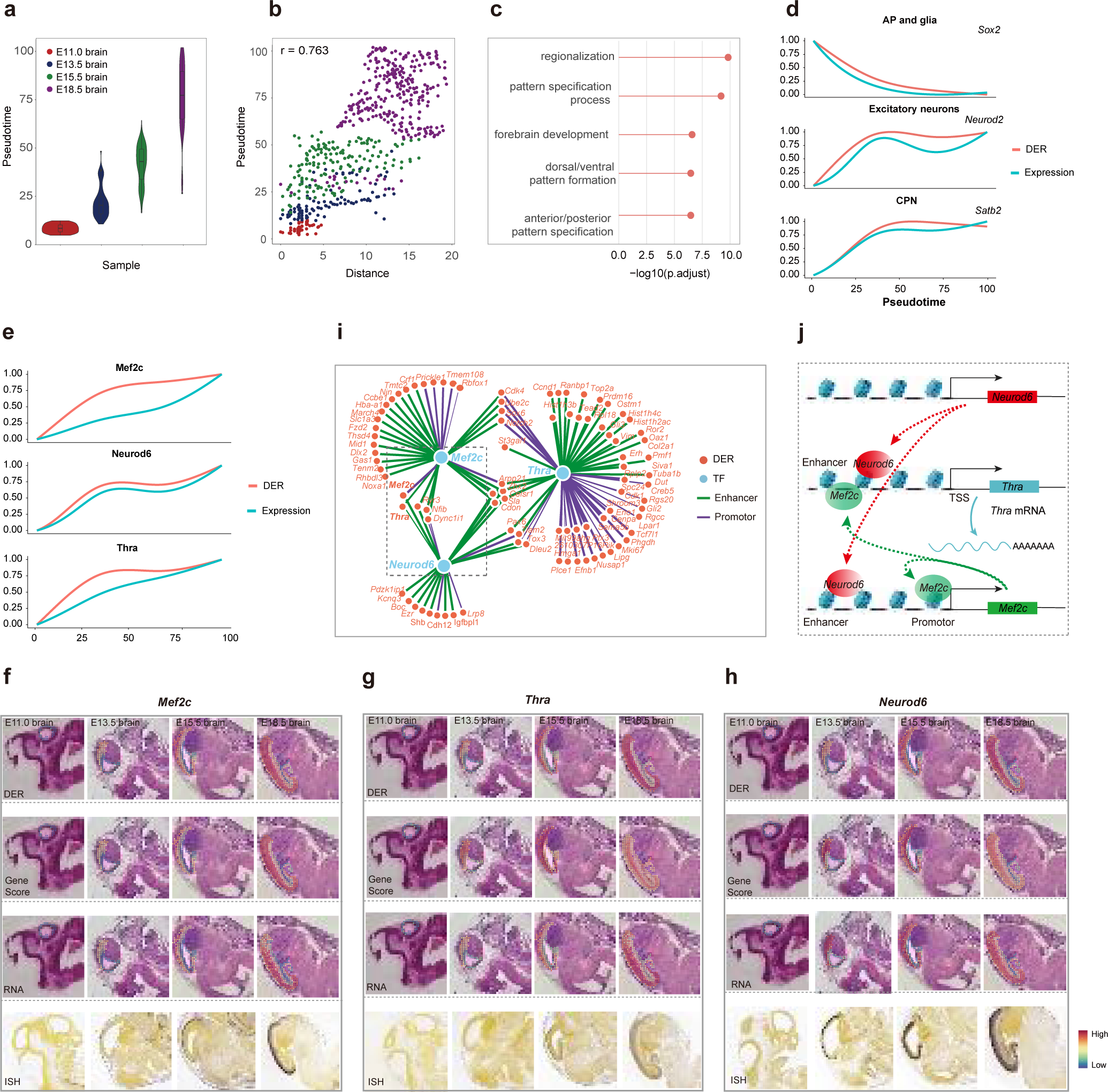

**Table S1. DNA oligos used for PCR and preparation of sequencing library.**

**Table S2. DNA barcode sequence.**

**Table S3. Stat of the MISAR-seq data.**

## Methods

### Mice

Mice were maintained in an animal facility approved by the Association for Assessment and Accreditation of Laboratory Animal Care (AAALAC), Guangzhou Institutes of Biomedicine and Health, Chinese Academy of Sciences. Procedures were approved by the Institutional Animal Care and Use Committees of GIBH (Institutional Animal Welfare Assurance Numbers N2019056). Normal brains were collected from wild-type mice of C57BL/6 mice at embryonic days 13.5-18.5. Mice of both sexes were used for the experiments.

### Mice brain cryosection

Mouse brains at different embryonic stages were dissected, washed with 1x PBS, embedded in OCT (Leica Microsystems), and stored at -80°C for tissue sectioning. Pre-cool the cryostat in advance, set the temperature to -20°C, and set the cutting thickness to 10 µm. Frozen blocks of OCT compounds were removed from cryo-embedding cassettes, and mouse brains from embryos at different stages were cut in sagittal planes, trimmed as needed, and attached to the metal stage of the OCT for cryosectioning. A piece of tissue is attached to each glass slice (188105, CITOTEST Scientific) and stored at -80°C.

### H&E Staining

The adjacent frozen slide was warmed at room temperature for 10 min and fixed with 200 μL 4% formaldehyde for 10 min. After decanting the formaldehyde, 200 μL isopropanol was added to the slide and incubate for 2 min. After completely dried, the tissue section was stained with 200 μL hematoxylin (sigma) for 4 min and cleaned in distilled water. The section was then incubated in 200 μL bluing reagent (sigma) for 1 min and rinsed in distilled water. At last, the section was stained with 200 μL eosin (sigma) for 3 min and cleaned in distilled water.

### Fabrication and assembly of microfluidic device

The device of microfluidic were fabricated using photo lithography. The thickness of the resistant and the channel width was 50 μm. The protocol of SU-8 photo lithography, development, and the hard baking was followed based on the manufacturer’s (MicroChem) recommendations. The fabricated molds of SU-8 were used to reprint microfluidic resin masters. Last, for microfluidic chips fabrication, polydimethylsiloxane (PDMS) was prepared by mixing base and curing agent at a 10:1 ratio and then poured to the resin masters. After degassing, the PDMS was cured (80°C, 2 h). PDMS slab was cut out after solidification and the inlet and outlet holes were punched for further use.

### Transposome preparation

Unmodified Tn5 was purchased from manufacturer Vazyme (S601-01), and the transposome was assembled following manufacturer’s guidelines. The oligos used for transposome assembly were as follows:

Tn5 Blocked_ME: /5Phos/C*T*G* T*C*T* C*T*T* A*T*A* C*A*/3ddC/

Tn5 Read 1: TCGTCGGCAGCGTCAGATGTGTATAAGAGACAG

Tn5 Pho_Read2: /5Phos/GTCTCGTGGGCTCGGAGATGTGTATAAGAGACAG

To evaluated the activity of the homemade Tn5, length distribution assay was performed using Qsep100 Bioanalyzer (Bioptic).

### Preparing oligonucleotides for ligations

There were two rounds of barcode chip ligation in MISAR-seq, each round using a different 50 barcodes (Table S2). Hybridization oligos have a universal linker sequence that is partially complementary to specific barcode sequences. These strands anneal before organizing barcodes to generate DNA molecules with three distinct functional domains: 5’ overhangs complementary to 5’ overhangs on cDNA molecules or transposition DNA molecules (may be derived from RT primers, transposition adapters, or a previous round of barcodes), a unique tissue-specific barcode sequence and another 5’ overhang that is complementary to the overhang on the DNA molecule to be ligated. To make 50 different round1 barcode mix in 96-well plate: add 25 μL 100 μM round 1 linker and 25 μL 50 different 100 μM barcodes to 50 different well and anneal the oligonucleotides by heating the plate to 95°C for 2 minutes and cooling to 20°C at a rate of about 1°C per minute. All oligonucleotides were dissolved in RNase-free water. The 50 different round2 barcode mix was prepared in the same way.

### Fixation and blocking

Sagittal sections of mouse brains at different embryonic stages, freshly cut or cryopreserved, were equilibrated at room temperature for 10 min. The tissue was fixed with 1% formaldehyde (252549-500ML, Sigma) for 5 minutes and quenched with 125 mM glycine for 5 min at room temperature. After fixation, the tissue section was quickly dipped in 1X PBS. Then, 1% BSA containing 0.25% RNase inhibitor was added to the tissue slide for blocking at room temperature for 30 min. The tissue was then ready for transposition.

### Transposition

The tissue section was permeabilized by Omni-ATAC buffer (10 mM Tris-HCl, pH 7.4; 10 mM NaCl; 3 mM MgCl2; 0.1%Tween-20 (P9416-50ML, Sigma); 0.1%NP-40 (11332473001, Roche); 0.01% Digitonin (D141-500MG, Sigma); 1%PIC (05892970001, Roche)) plus 0.25% RNase inhibitor for 10 minutes at room temperature, then was washed with ATAC-RSB buffer (10 mM Tris-HCl pH 7.4; 10 mM NaCl; 3 mM MgCl2; 0.1% Tween-20; 0.25% RNase inhibitor) at room temperature for 5 min. And then the tissue section was washed by RNase free water for three times and air dried. 50 µL transposition buffer containing 0.25% RNase inhibitor (25 µL 2X TD buffer (20% Dimethylformamide; 20 mM Tris-HCl pH 7.4; 10 mM MgCl2); 19.4 µL 1X DPBS; 0.5 µL 10% Tween-20; 0.1 µL 5% Digitonin; 5 µL transposome) was added followed by incubation at 37°C for 30 minutes. To stop tagmentation, 100 µL TS buffer (20 mM EDTA; 1 mM Spermidine) containing 0.25% RNase inhibitor were added after removing transposition buffer. The tissue slide was washed with ATAC-RSB buffer for 1 minute. And then the liquid was removed, the tissue slide was fixed with 4% formaldehyde at room temperature for 20 min and washed with ATAC-RSB buffer and 0.5x PBS.

### Reverse transcription

Place tissue sections in a humidified box, and each tissue section was mixed with 45 μL of RT mix (final concentration of 1x HiScript III buffer, 0.33 M/μL RNase Inhibitor, 0.5x PBS, 20 M/μL HiScript III Reverse Transcriptase (R302, Vazyme), 500 μM dNTP (18427088, Thermo) and 3.5 μM RT primer with an affinity tag). The RT primer contains a poly-T tail, a Unique Molecular Identifier (UMI), a universal ligation overhang, and a biotin molecule. The RT reaction was done with regular procedure.

### Ligation

200 μL of ATAC-RSB buffer containing 0.25% RNase inhibitor were added to the tissue area. The slide was gently washed by 200 μL of 1x NEBuffer 3.1. For the first round of ligation, longitudinal PDMS chips were attached in the area of interest and PDMS device were then clamped with an acrylic clamp. The brightfield image was taken with 4x objective (OLYMPUS microscope) for further alignment. 3 μL of Round1 barcode mix (1x T4 DNA ligase buffer, 16 M/μL T4 DNA ligase (M0202L, NEB), 0.25 M/μL RNase Inhibitor, NF-H2O, 1x NEBuffer 3.1, 5 μM Cy3/Fam dye, 50 μM round1 barcode mix) was added to each inlet well of the chip. The fluid was pulled over slowly with a syringe and the reaction was carried out at 37°C, 80 rpm for 30 min in a wet box. After the reaction, the Cy3 and Fam images were taken by fluorescence photography. After discarded the ligation solution, the clamp and PDMS chip were removed and the tissue slide was washed by RNase free water. 200 μL of ATAC-RSB buffer was added to the tissue area. The second round of barcode ligation was similar to the first round.

### Tissue digestion

50 μL 2x Lysis buffer (20 mM Tris, pH8.0; 400 mM NaCl; 100 mM EDTA, pH8.0; 4.4%SDS; NF-H2O) and 10 μL proteinase K (20 mg/mL) (RT403-02, TIANGEN) was added to the slide and incubated for 10 min at room temperature. The lysate was further digested at 55 °C 2 h and collected into a 1.5 mL EP tube.

### cDNA library preparation

RNA sequencing library was generated following the SHARE-seq protocol as previously reported with slight modifications. Briefly, the qPCR reaction was carried out at the following conditions: 95°C for 3 min, and then 30 thermal cycles at 98°C for 20 s, 67°C for 20 s and 72°C for 3 min. The PCR products were purified by 0.7X Ampure XP beads (A63882, BECKMAN COULTER) and dissolved in 15 μL RNase free water. cDNA concentration and fragment size distribution were determined by Qsep100 (Bioptic).

### MISAR-seq RNA library preparation

The cDNA library construction was performed according to the manufacturer’s guidelines of TruePrep DNA Library Prep Kit V2 for Illumina (TD502, Vazyme).

### MISAR-seq ATAC library preparation

Zymo DNA Clean & Concentrator-5 (D4014, ZYMO research) was used to extract the genomic DNA in the supernatant retained after C1 beads purification, and eluted with 25 μL of NF-H2O. 20 μL of the eluted purified product was then mixed with the PCR solution (2.5 µL 10 µM Ad1.01∼Ad1.08 primer; 2.5 µL 10 µM P7 primer; 25 µL 2x NEBNext Master Mix (M0541, NEB)). Then, PCR was conducted with following the program: 58°C for 5 min, 72 °C for 5 min, 98 °C for 30 s, and then cycled 5 times at 98 °C for 10 s, 65°C for 10 s, and 72°C for 1 min. To determine additional cycles, 5 µL of the pre-amplified mixture was first mixed with the qPCR solution (0.5 µL10 µM Ad1.01∼Ad1.08 primer; 0.5 µL 10 µM P7 primer; 1 µL 20x SYBR Green (31000, BIOTIUM); 10 µL 2x NEBNext Master Mix; 2 µL NF-H2O). Then, qPCR reaction was carried out at the following conditions: 98°C for 30 s, and then 30 cycles at 98°C for 10 s, 65°C for 10 s, and 72°C for 1 min. Finally, the remainder 45 µL of the pre-amplified DNA was amplified by running the required number of additional cycles of PCR (cycles needed to reach 1/3 of saturated signal in qPCR). The PCR products were further purified by 1X Ampure XP beads and dissolved in 15 μL RNase free water.

### Sequencing strategy

Both libraries were sequenced on an MGI-seq 2000 sequencer (BGI company) using a pairend mode. cDNA library was sequenced with read1 75 bp, read2 10 bp, index5 8 bp and index7 50 bp. ATAC library was sequenced with read1 75 bp, read2 75 bp, index5 8 bp and index7 50 bp.

### MISAR-seq data preprocessing

The sequences were transformed to Cell Ranger ARC format using a custom python script which modified from spatial-ATAC-seq. Resulting fastq files were aligned to mouse genome (mm10) and counted using Cell Ranger ARC v2.0.1. The fragment file and feature barcode matrix file were generated for downstream analysis. To filter the data in non-tissue regions, microscope fluoresce images with two channels were used to identify the pixel locations with manual selection using Adobe Photoshop and generated a mask picture which tissue region was filled with white color while non-tissue region was filled with black color. Then we can use a custom python script to get the filtered barcode location and used to filter fragment file and feature barcode matrix file. The filtered fragments file was then read into ArchR as a tile matrix in 5kb genome binning size using createArrowFiles and ArchRProject function. The filtered feature barcode matrix file was also read into the same ArchR project using import10xFea tureMatrix and addGeneExpressionMatrix function. Besides, the barcode spatial position matrix was read into the same ArchR project using addCellColData function. Both ATAC and RNA data normalization and dimensionality reduction was performed using Latent Semantic Indexing (LSI) and batch effects was removed by addHarmony function. Besides, the combined of ATAC and RNA reduced dimensions was generated from addCombinedDims function using ATAC and RNA batch removed reduced dimensions. Then, all graph clustering (addClusters function) and Uniform Manifold Approximation and Projection (UMAP) embeddings (addUMAP function) using the batch removed reduced dimensions with default parameters. Gene Score matrix was generated using Gene Score model. Cluster marker genes were calculated using the getMarkerFeatures and getMarkers function (testMethod = "wilcoxon", cutOff = "FDR <= 0.05 & log2FC >= 1"). Using the MACS2 to call peaks and motif enrichment or deviations were calculated using peakAnnoEnrichment and addDeviationsMatrix function. Using the clusterProfiler package to perform GO enrichment analysis (qvalueCutoff = 0.05). To visualize data, the clusters or variable genes were projected into 2D space by UMAP using plotEmbedding function. Spatial clusters were plotted to adjacent H&E-stained tissue by custom R script based on ggplo2 according to the relationship of barcode and location (round1 barcode as column and round2 barcode as row). All functions used in above are from ArchR package.

### Genes and umi comparison

To equitably compare the number of genes and umis with or without 4% refix before reverse transcription, we use preseq software to exact the same 10M distinct reads for all of the four samples and then performed the barcode split and gene count analysis as MISAR-seq data preprocessing described.

### The integration of single cell RNA-seq data and MISAR-seq data

We retrieved scRNA data from the developing mouse brain^2^. We downloaded the loom file from Mouse Brain Atlas (http://mousebrain.org/development/downloads.html), which including all gene expression values and annotated metadata per cell. Then the reference data was converted into a seurat object using the scanpy python package of scanpy and R package of Seurat. The E15.5 brain data was selected and the pseudotissue cell type size < 25 was filtered out. Meanwhile, mitochondrial and hemoglobin gene also removed for downstream analysis. For E15.5 brain MISAR-seq data, we first read the feature barcode matrix into seurat use Read10X_h5 function from Seurat. The Seurat RNA and ATAC assay was generated by CreateSeuratObject function from Seurat and CreateChromatinAssay function from Signac. Then, for RNA, the SCT assay generated from RNA assay using SCTransform function in Seurat with do.scale = T. For ATAC, the peaks assay was created from new peaks matrix generated by macs2 using the CallPeaks function in Signac. The gene activities assay was generated by GeneActivity function in Signac. For integration of MISAR-seq RNA data with reference, we first use FindIntegrationAnchors function in Seurat of top 2000 high variable genes to find their anchors and then use IntegrateData function in Seurat to integrate both of them. At last, the new generated integrated assay was used to perform dimensionality reduction and graph clustering use functions in Seurat. The reference data label of pseudotissue was plot with DimPlot function in Seurat. For integration of MISAR-seq ATAC data with reference, we first use the activity assay to perform FindTransferAnchors function in Seurat to find their anchors and then use TransferData function in Seurat to get a imputed scRNA-seq matrix for each of the ATAC grids. After merge reference data and MISAR-seq ATAC data, the new generated combined object was used to perform dimensionality reduction and graph clustering use functions in Seurat. The reference data label of pseudotissue was plot with DimPlot function in Seurat; https://satijalab.org/seurat/index.html.

### The single cell deconvolution of MISAR-seq RNA data

The reference of single cell data and MISAR-seq RNA data is almost the same as used in integration above except for the cell type of subclass. For cell-type deconvolution, we used RCTD to deconvolve cell types within a given grid after filtered out cell type size < 25, mitochondrial and hemoglobin gene. We used create.RCTD and run.RCTD function in spacexr package with the ‘multi’ mode and a mini-mum of 25 cells for each cell-type identity to deconvolve. Other parameters were used with the default; https://github.com/dmcable/spacexr.

### Peak co-accessibility and peak-gene linkages

To calculate the peak co-accessibility, we use addCoAccessibility function and getCoAccessibility function from ArchR with default parameters. To find the significant peak-gene linkages, we use addPeak2GeneLinks function and getPeak2GeneLinks function from ArchR with default parameters excepted useMatrix = “GeneExpressionMatrix”. To find the *Neurod1* and *Rfx2* motifs in significant peak gene linkages, we exacted the motif position weight matrix (pwm) from chromVARmotifs package and use matchMotifs function from motifmatchr package to match motifs site in peaks. To plot both co-accessibility and peak-gene linkages in genome track, we use plotBrowserTrack function in ArchR with default parameters excepted useMatrix = "GeneExpressionMatrix".

### Define spatiotemporal development trajectory of mouse cortex

To incorporate gene expression and peak accessibility to infer corticogenesis trajectory, we selected two clusters that aligned well with cortex anatomy structure and concatenated two batch-removed harmony embedding matrixes as combined latent space. The cellular distance was calculated based on this combined latent space and then fed into the supervised trajectory inference method implemented in the ArchR package.

### Linking ATAC peaks and gene expression in the trajectory axis of mouse cortex

To connect cis-regulatory elements with target gene expression, we computed the rolling mean average of peak accessibility and gene expression across the corticogenesis trajectory axis and calculated the Pearson correlation coefficient based on smooth value. Only peaks within 250kb on either side of TSS were considered for each gene, especially excluded promoter regions. We retained peak-gene links with FDR < 0.01 and PCC > 0.1, which resulted in 66805 significantly putative enhancer-gene linkages. Inspired by the conception of the domain of region accessibility (DORC), we defined domain of enhancer regulatory (DER) by aggregating all linked peak counts of each target gene in a distance-dependent fashion. A generalized linear model was performed to test each gene and accessibility activity for differential expression as a function of pseudotime, which resulted in 1160 putative enhancer-gene interaction pairs. Those interaction pairs were clustered into 5 modules using hierarchical clustering based on euclidean distance. Enrichment analysis was performed using the ClusterProfier for each gene module. The top 5 significant GO terms for each module were shown (FDR < 0.01).

### Gene regulatory network analysis

Accessibility activity score and gene expression counts were smoothed using 30 nearest neighbors in combined latent space and then fed together with peak-gene links to FigR function ’runFigRGRN’ to construct gene regulatory network. TF ranked by the mean regulation score with function ’rankDrivers’.

### Data quality comparison with other methods

E15.5 mouse forebrain bulk ATAC-seq data was downloaded from ENCODE and E11.0 whole mouse DBiT-seq data (50 μm resolution) was downloaded from GSE137986. Besides, in situ hybridization and expression images were downloaded from Allen Mouse Brain Atlas.

## Notes

### Competing Interest Statement

The authors have declared no competing interest.

### Summary of Updates

Title was modified and new figures were added.

